# The P681H mutation in the Spike glycoprotein escapes IFITM restriction and is necessary for type I interferon resistance in the SARS-CoV-2 alpha variant

**DOI:** 10.1101/2022.08.11.503706

**Authors:** Maria Jose Lista, Helena Winstone, Harry D Wilson, Adam Dyer, Suzanne Pickering, Rui Pedro Galao, Giuditta De Lorenzo, Vanessa M. Cowton, Wilhelm Furnon, Nicolas Suarez, Richard Orton, Massimo Palmarini, Arvind H. Patel, Luke Snell, Gaia Nebbia, Chad Swanson, Stuart J D Neil

## Abstract

The appearance of new dominant variants of concern (VOCs) of severe acute respiratory syndrome coronavirus type 2 (SARS-CoV-2) threatens the global response to the COVID-19 pandemic. Of these, the alpha variant (also known as B.1.1.7) that appeared initially in the UK became the dominant variant in much of Europe and North America in the first half of 2021. The Spike (S) glycoprotein of alpha acquired seven mutations and two deletions compared to the ancestral virus, including the P681H mutation in the polybasic cleavage site that has been suggested to enhance S cleavage. Here, we show that the alpha S protein confers a level of resistance to the effects of interferon-β (IFNβ) in human lung epithelial cells. This correlates with resistance to an entry restriction mediated by interferon-induced transmembrane protein 2 (IFITM2) and a pronounced infection enhancement by IFITM3. Furthermore, the P681H mutation is essential for resistance to IFNβ and context-dependent resistance to IFITMs in the alpha S. However, while this appears to confer changes in sensitivity to endosomal protease inhibition consistent with enhanced cell-surface entry, its reversion does not reduce cleaved S incorporation into particles, indicating a role downstream of furin cleavage. Overall, we suggest that, in addition to adaptive immune escape, mutations associated with VOCs may well also confer replication and/or transmission advantage through adaptation to resist innate immune mechanisms.

**IMPORTANCE:** The emergence of Variants of Concern of SARS-CoV-2 has been a key challenge in the global response to the COVID-19 pandemic. Accumulating evidence suggests VOCs are being selected to evade the human immune response, with much interest focussed on mutations in the Spike protein that escape from neutralizing antibody responses. However, resistance to the innate immune response is essential for efficient viral replication and transmission. Here we show that the alpha (B.1.1.7) VOC of SARS-CoV-2 is substantially more resistant to type-1 interferons than the parental Wuhan-like virus. This correlates with resistance to the antiviral protein IFITM2, and enhancement by its paralogue IFITM3, that block virus entry into target cells. The key determinant of this is a proline to histidine change at position 681 in S adjacent to the furin-cleavage site that we have shown previously modulates IFITM2 sensitivity. Unlike other VOCs, in the context of the alpha spike, P681H modulates cell entry pathways of SARS-CoV-2, further reducing its dependence one endosomal proteases. Reversion of position 681 to a proline in viruses bearing the alpha spike is sufficient to restore interferon and IFITM2 sensitivity without reducing furin-mediated spike cleavage, suggesting post cleavage conformational changes in S are changing the viral entry pathway and therefore sensitivity to interferon. These data highlight the dynamic nature of the SARS CoV-2 S as it adapts to both innate and adaptive immunity in the human population.

## INTRODUCTION

Both SARS-CoV-1 and SARS-CoV-2 enter target cells through the interaction of their S proteins with the angiotensin converting enzyme 2 (ACE2) cell surface receptor. Upon attachment and uptake, the S glycoprotein trimer is cleaved by cellular proteases such as cathepsins and TMPRSS family members at two positions – the S1/S2 junction and the S2’ site – to facilitate the activation of the fusion mechanism. Similar to more distantly related beta-CoVs, but so far unique in known Sarbecoviruses, the SARS-CoV-2 glycoprotein contains a polybasic furin cleavage site (FCS) with a (681-PRRAR*S-685) sequence at the S1/S2 junction. This allows the S precursor to be additionally processed to the S1 and S2 subunits by furin-like proteases before viral release from the previously infected cell [1]. This leads to a proportion of processed S to be present on the virion before engagement with the target cell, allowing for rapid activation and fusion at or near the cell surface by TMPRSS2. The importance of the FCS is highlighted by the observations that it enhances SARS-CoV-2 replication specifically in airway epithelial cells and is essential for efficient transmission in animal models [2].

The alpha variant of SARS-CoV-2 arose in the South-East of England in autumn 2020, and rapidly spread across the world in the first months of 2021. Various studies suggested that alpha had an increased transmissibility between individuals [3-5]. Alpha contains nine amino acid residue changes in S including a deletion of amino acid residues H and V in the N-terminal domain at position 69/70, thought to increase S incorporation into virions, a single amino acid deletion of Y144 (thought to assist NTD antibody neutralization escape), and a N501Y mutation in the RBD which enhances ACE2 binding affinity [6, 7]. Together these changes have been shown to reduce efficiency of neutralization by some antibodies [8] but compared to the later VOCs Delta and Omicron, it is not thought to be a major adaptive immune escape variant. Alpha also acquired a P681H change in the FCS which has been proposed to increase the accessibility of the site by furin leading to enhanced cleavage as well as more efficient cell-to-cell fusion and syncytia formation [9-12]. Since early 2021, several other VOCs have emerged with mutations in the FCS, including kappa, delta, and omicron [12, 13]. Both kappa and delta contained the P681R mutation, however only delta superseded alpha and became a globally dominant variant in the summer of 2021. In late 2021, the delta variant was then in turn displaced by the omicron variant, which contains the P681H mutation in its FCS.

We and others have previously demonstrated that the ancestral SARS-CoV-2 is variably sensitive to entry inhibition by the interferon-regulated IFITM family and that this can be modulated by the FCS [2, 14, 15]. IFITMs 1, 2 and 3 are transmembrane proteins that exert antiviral activity against diverse enveloped viruses by blocking fusion of the viral and cellular membranes [16, 17]. While IFITM1 localizes primarily to the plasma membrane, IFITM2 and IFITM3 are internalized via a conserved YxxΦ endocytic motif to occupy distinct and overlapping endosomal compartments. However, it has previously been demonstrated that the IFITM proteins can oligomerize with each other in heterologous complexes [18, 19]. The sensitivity of a given virus to individual IFITM proteins is largely determined by their route of cellular entry. We have previously shown that for a prototypic Wuhan-like SARS-CoV-2 isolate from early 2020, IFITM2 reduced viral entry and contributed to type I interferon (IFN-I)-induced inhibition in human cells [14]. Sensitivity to IFITM2 could be markedly enhanced by deletion of the FCS, suggesting that furin processing ameliorated SARS-CoV-2 sensitivity to IFITM2-restriction at least to some extent. We therefore postulated that the altered cleavage site of VOCs with mutations in the FCS may have consequences for their sensitivity to IFN-I and IFITMs. Here, we demonstrate that of the alpha, beta, gamma, kappa, delta and omicron variants, only the S of the alpha variant is resistant to IFITM restriction in A549-ACE2-IFITM cells. We also demonstrate that the ΔCT mutation commonly used in improving SARS-CoV-2 PLV infectivity masks the IFITM resistance of alpha PLVs by conferring increased cathepsin-dependence. Furthermore, we show that the alpha variant is resistant to IFNβ in both A549-ACE2 and Calu-3 cells, which can be abolished by reversion of the P681H mutation.

## RESULTS

### The S proteins of currently circulating variants display differing sensitivities to IFITMs in A549-ACE2 cells

Previously we have shown that viral entry mediated by the original Wuhan-1 S pseudotyped lentiviral vector (PLV) or the England-02 isolate (hCoV-19/England/02/2020) was inhibited by IFITM2 in A549-ACE2 cells, and that this effect correlated in part with the IFNβ sensitivity of the virus [14]. Over 2020 and 2021 several major variants of concern (VOCs) have arisen – alpha (B.1.1.7) in the UK, beta (B.1.351) in South Africa, gamma (P1) in Brazil, delta (B.1.617.2) in India, and most recently, the omicron family (B.1.1529) in South Africa [13]. All of these variants have multiple changes in the S protein that could potentially affect the entry process (Figure 1). Of particular interest, the alpha, delta and omicron variants contain mutations in the polybasic cleavage site which have been postulated to enhance S cleavage: P681H in alpha and omicron, and P681R in delta [20-22]. We therefore compared the sensitivity of full-length S PLV entry of these VOCs in the presence of IFITM proteins. As expected, all VOC PLVs produced were infectious on A549-ACE2 cells, although efficiency was variable (Supplementary Figure 1A). We then used these PLVs to infect A549-ACE2 cells stably expressing the individual IFITMs (Supplementary Figure 1B, Figure 2A-2I). The D614G mutation that became dominant early in the first wave of the pandemic displayed a similar sensitivity to IFITM2 as the previously characterized Wuhan-1 S, but was resistant to both IFITM1 and IFITM3 (Figure 2A, 2B) [14, 23]. We then compared the IFITM sensitivities of alpha, beta, gamma, kappa, delta, omicron (BA.1), and omicron (BA.2) as PLVs (Figure 2C-2I).The alpha S (Figure 2C) appeared completely insensitive to IFITMs 1, 2 or 3 whilst beta, gamma, kappa, delta, and both omicron spikes still retain some sensitivity to IFITMs 1 and/or 2. We noted that kappa and delta (Figure 2F, 2G), which both contain the P681R mutation, retained some sensitivity to both IFITMs 1 and 2. Interestingly the alpha variant, and to some extent delta, also appeared to be significantly enhanced by IFITM3. Such enhancement by IFITMs has been previously documented in the human seasonal CoV OC43, and in SARS-CoV-2 under specific circumstances [24, 25]. To confirm the enhancement we observed with alpha was due to IFITM3, we pre-treated A549-ACE2-IFITM3 cells with cyclosporin H, a compound known to drive IFITM3 to ubiquitin-dependent degradation and found that this led to specific abolishment of IFITM3 enhancement of alpha PLVs while having no effect on D614G (Supplementary Figure 1C, 1D) [9, 26][9, 25, 26].

**Figure 1.**
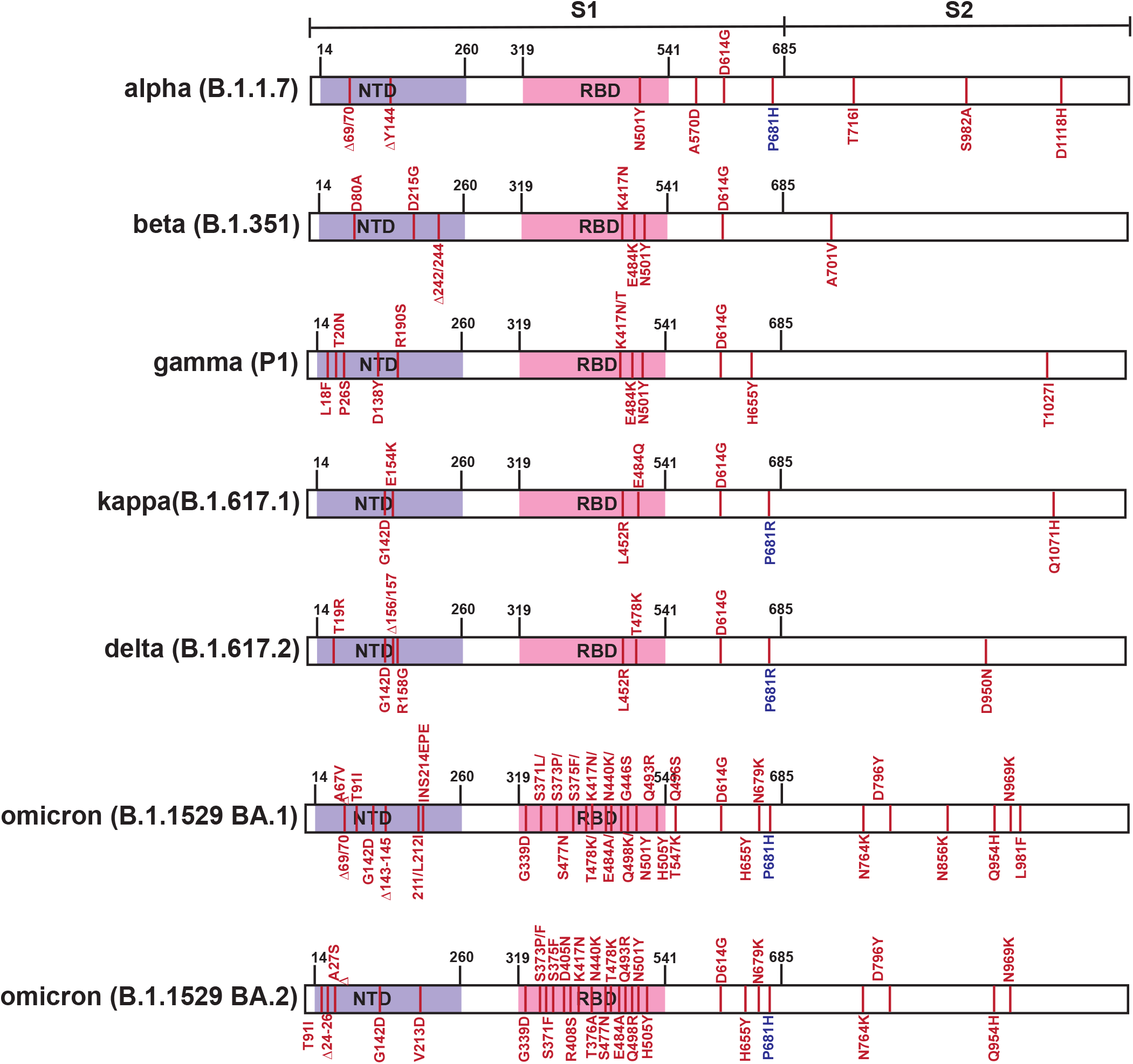
SARS-CoV-2 variants of concern spike sequences. Schematic of Spike protein domains of the different variants of concern relative to the original Wuhan Spike sequence: alpha, beta, gamma, delta and omicron. The different mutations between the variants are represented in red.

**Figure 2.**
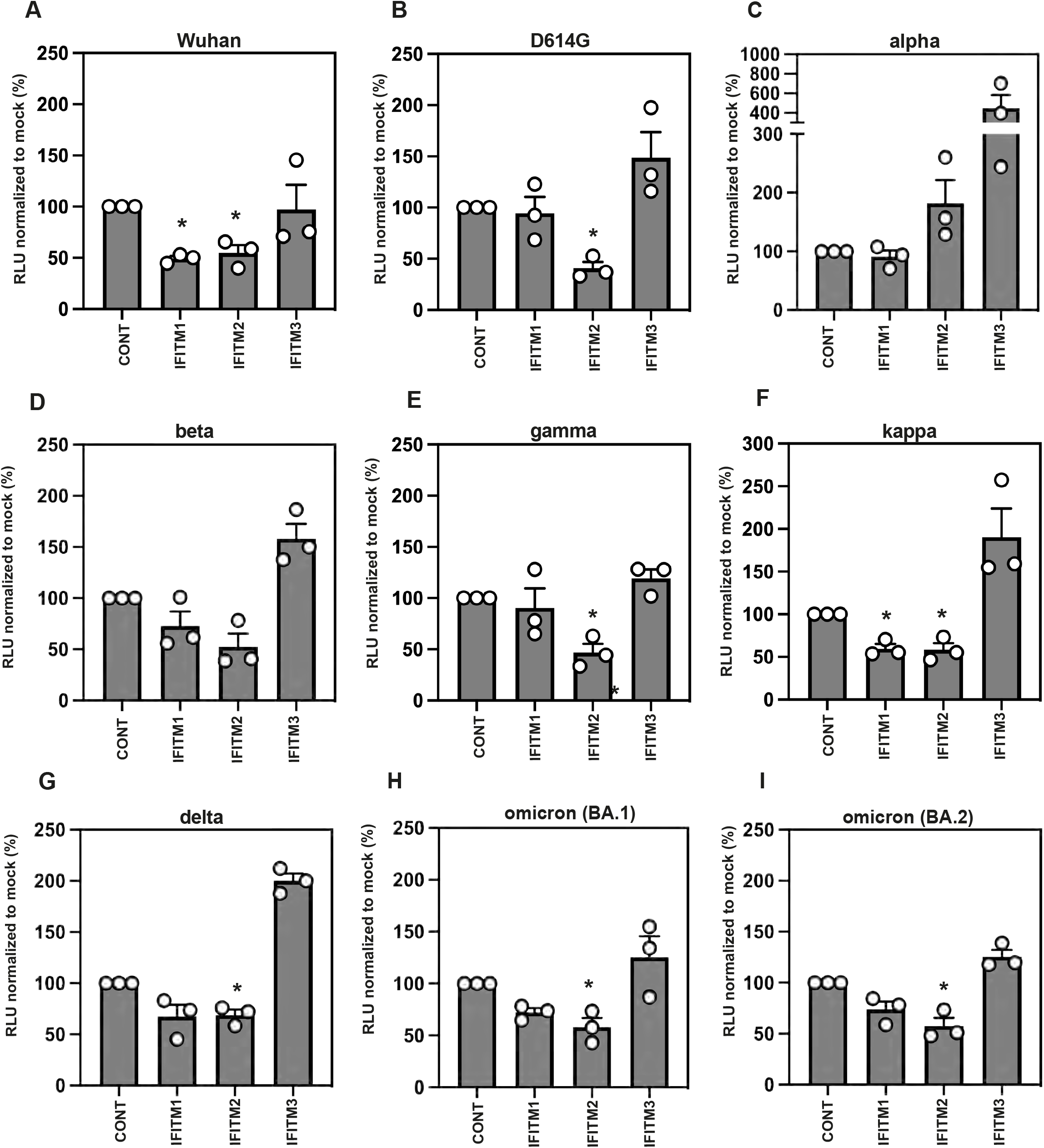
IFITM sensitivity of SARS-CoV-2 variants of concern. A-I) IFITM sensitivity of Wuhan, D614G, alpha, beta, gamma, kappa, delta and omicron PLVs in A549-ACE2 cells stably expressing the individual IFITMs. PLV entry was quantified by Luciferase activity 48 hours after infection and normalized to control cells. Data shown are mean ± SEM, n=3. Statistics were calculated in Prism using *ANOVA*, stars indicate significance between control cell and individual IFITM (*P=0.05).

### The ΔCT mutation increases PLV infectivity but confers greater cathepsin-dependence and IFITM2 sensitivity to D614G and alpha PLVs

Deleting the last 19 amino acids of SARS-CoV-2 spike increases spike incorporation and infectivity of PLVs and is common practice amongst many groups studying SARS CoV-2 [27, 28]. Truncation of the cytoplasmic tail results in the deletion of a sub-optimal endoplasmic reticulum retention signal (ERRS) and increased accumulation of the spike at the surface where it is incorporated into PLVs. However, the site of coronavirus assembly is not at the plasma membrane and the spike goes through considerable post-translational modifications in the ERGIC [29]. To test whether deletion of the last 19 amino acids affected IFITM phenotypes, we generated a D614GβCT mutant and tested infectivity in A549-ACE2 cells of these PLVs relative to the full-length D614G spike as PLVs (Figure 3A). The ΔCT mutant exhibited a 28-fold boost in infectivity (Figure 3A). However, the D614GΔCT PLVs were 2-fold more sensitive to IFITM2 (Figure 3B). This was consistent with an increase in sensitivity of these PLVs to E64d, an inhibitor of cathepsins B/L at both 2.5*μ*,M and 10*μ*M (Figure 3C). Next, to confirm if there were phenotypic differences in the spike of D614GΔCT spikes during PLV production, D614G and D614GΔCT were immunoblotted for Spike and Gag in both the cell lysates and purified supernatant of PLV production (Figure 3D). Intriguingly, the D614GΔCT mutant showed increased S1/S2 processing by 10-fold (Figure 3E). Although increased spike processing was surprising given an increased dependence on cathepsins B/L, it could be that although more processed, the D614GΔCT spike is in a conformation such where the second cleavage site is less accessible, resulting in creased cathepsin-dependence. Finally, to confirm whether the ΔCT mutation is sufficient to overcome the IFITM2 resistance observed with the alpha spike, alphaΔCT was generated and the IFITM sensitivity of this tested (Figure 3F). Strikingly, the ΔCT mutation rendered the previously resistant alpha spike highly sensitive to IFITM2. Additionally, the 3-fold enhancement we previously found with alpha in this system was abolished by the ΔCT mutation. Overall these data suggest that the ERRS plays a significant role in the post-translational modifications of spike and in turn, this has consequences for the route of viral entry and sensitivity to antiviral proteins. Given the significant effect of this mutation on IFITM sensitivity and route of viral entry of D614G and alpha, we would advise caution from interpreting data of phenotypes involving differential viral entry utilising ΔCT spikes.

**Figure 3.**
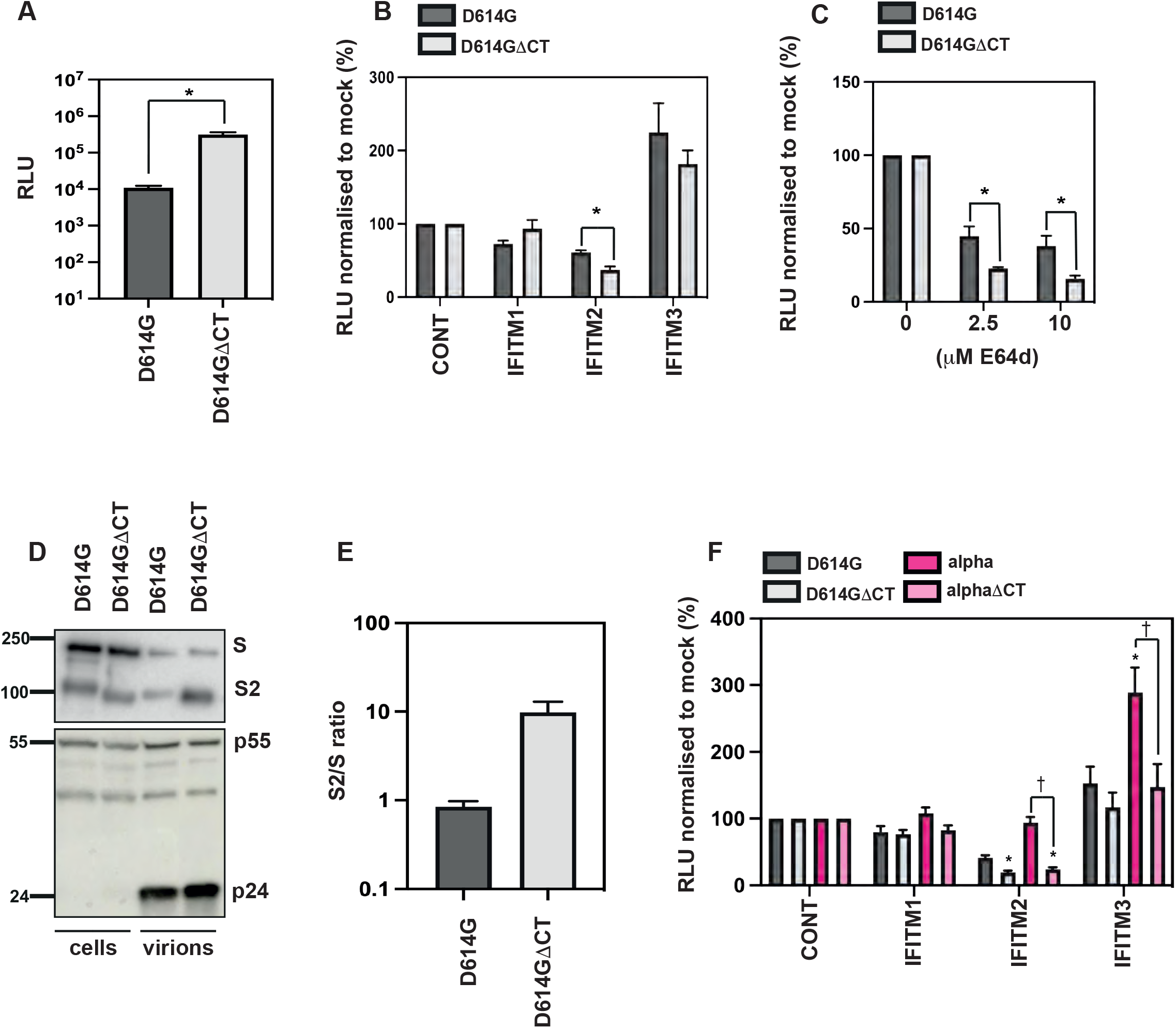
The ΔCT mutation in D614G or alpha confers IFITM2 sensitivity by increasing cathepsin-dependence.A) D614G or D614GΔCT PLVs were used to infect A549-ACE2 cells and infectivity measured by Luciferase activity 48 hours later. Raw RLUs shown. B) D614G or D614GΔCT PLVs were used to infect A549-ACE2-IFITM cells and infectivity measured by Luciferase activity 48 hours later. Percent infection normalised to control cells without IFITM shown. C). A549-ACE2 cells were pre-treated with 2.5*μ*M or 10*μ*M E64d prior to infection with D614G or D614GΔCT PLVs for 48 hours. Infection was measured by Luciferase activity and infection normalised to mock treated cells. D) Representative immunoblot of cell lysates and supernatant from PLV production. Supernatant was purified through a 20% sucrose cushion for 1 hour at 18000 G prior to lysis. E) Quantification of 3 independent immunoblots of ratio of S2 over S of purified supernatant of D. F) D614G, ΔCT, alpha or alpha ΔCT PLVs were used to infect A549-ACE2-IFITM cells for 48 hours and infection quantified by Luciferase activity. Infection is normalised to control cells. Data shown are mean ± SEM, n=3. Statistics were calculated in Prism using ANOVA, stars indicate significance between control cell and individual IFITM and crosses indicate significance between different IFITM/drug conditions (P=<0.05).

### SARS CoV-2 alpha variant is IFITM resistant

Next, we sought to confirm that the native alpha virus demonstrated a similar phenotype on our IFITM-expressing cells as the PLVs. We infected A549-ACE2 cells stably expressing the individual IFITMs with England-02, D614G, or alpha isolates and measured the percentage of N positive cells by flow cytometry (Figure 4A, 4B), and intracellular E RNA by qPCR (Figure 4C) at 48h post-infection. We found that England-02 and D614G were IFITM2 sensitive, while alpha was insensitive to the effects of all three IFITMs. We also noted again significant enhancement of infection in the presence of IFITM3, and to a lesser extent IFITM1, consistent with our PLV experiments. Furthermore, both delta and omicron viruses displayed sensitivity to both IFITM2 and IFITM3 (Supplementary Figure 2). Thus, the alpha variant of SARS CoV-2, unique amongst the current VOCs, is fully IFITM resistant in A549-ACE2s. Furthermore, the IFITM3 enhancement of alpha infection is reproducible between PLVs and native virus.

**Figure 4.**
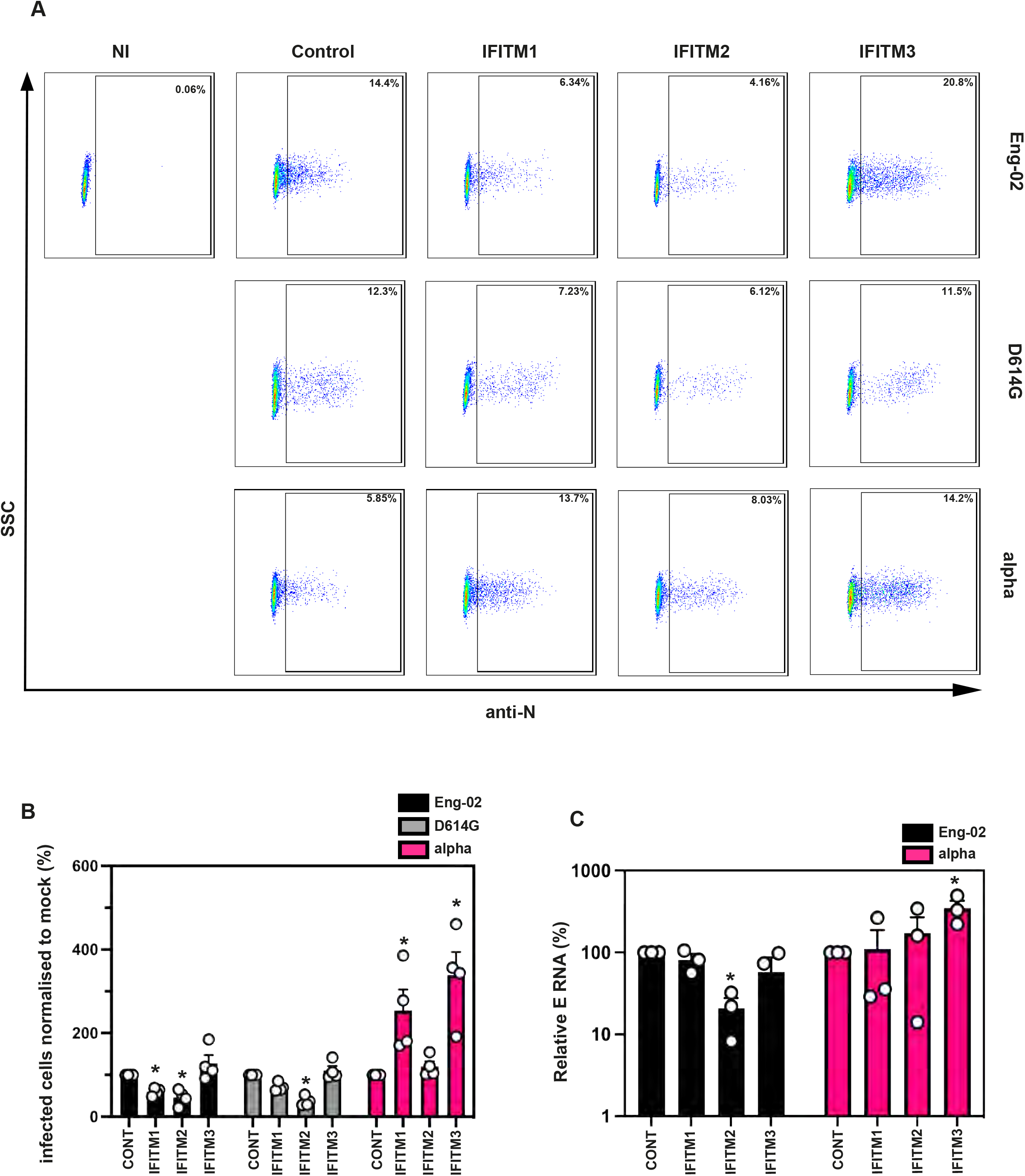
The alpha variant of SARS-CoV-2 is resistant to IFITMs. A) representative FACS plots of intracellular N staining of infected A549-ACE2-IFITM cells. NI= non-infected. B) Quantification of intracellular N staining by flow cytometry of A549-ACE2 IFITM cells infected with England 02, Wuhan D614G, and alpha. A549-ACE2 expressing the individual IFITMs were infected with isolates of England 02, D614G or alpha for 48h. Infection was measured by percentage of N positive cells by flow cytometry. Data analysed in FlowJo. C) Infection of A549-ACE2 stably expressing the individual IFITMs with England 02 and alpha viruses at MOI 0.01. Infection was quantified by RT-qPCR of E gene relative to GAPDH 48 hours later; graph represents E mRNA levels relative to GAPDH. Data shown are mean ± SEM, n=3. Statistics were calculated in Prism using ANOVA, stars indicate significance between control cell and individual IFITM and crosses indicate significance between different IFITM/drug conditions (P=<0.05).

### The alpha variant is less sensitive to IFNβ than an early pandemic isolate

While previous data has indicated that the original Wuhan-like SARS-CoV-2 virus can delay pattern recognition of viral RNA in target cells, its replication is highly sensitive to exogenous IFN-I treatment in culture, in part determined by IFITM2 [30]. Having confirmed that the alpha variant is resistant to IFITM expression when ectopically expressed, we then tested if alpha was also more resistant to the effects of IFNβ, as suggested by others [31, 32]. Indeed, we found from measuring supernatant viral RNA 48 hours post-infection of A549-ACE2 cells that alpha is more resistant than England-02 to pre-treatment with increasing doses of IFNβ (Figure 5A). Additionally, this was recapitulated in the Calu-3 cell line, which endogenously express ACE2 and more faithfully represent target cells in the respiratory tract (Figure 5B). We further extended these observations to two clinical isolates of alpha (clinical isolates 10 and 28; Figure 5C) and measured viral RNA in cell lysates. This confirmed that two clinical isolates of alpha grown from patient swabs are also resistant to pre-treatment with IFNβ. Finally, we showed that the alpha isolate is resistant to exogenous IFNβ pre-treatment by taking the supernatant from infected Calu-3 cells pre-treated with IFNβ and measuring the viral infectivity by plaque assay on Vero-E6-TMPRSS2 cells, confirming that the alpha variant still actively replicates in the presence of IFNβ to produce infectious virions (Figure 5D). Thus, in comparison to a representative example of Wuhan-1-like SARS-CoV-2, the alpha variant has a marked resistance to IFN-I.

**Figure 5.**
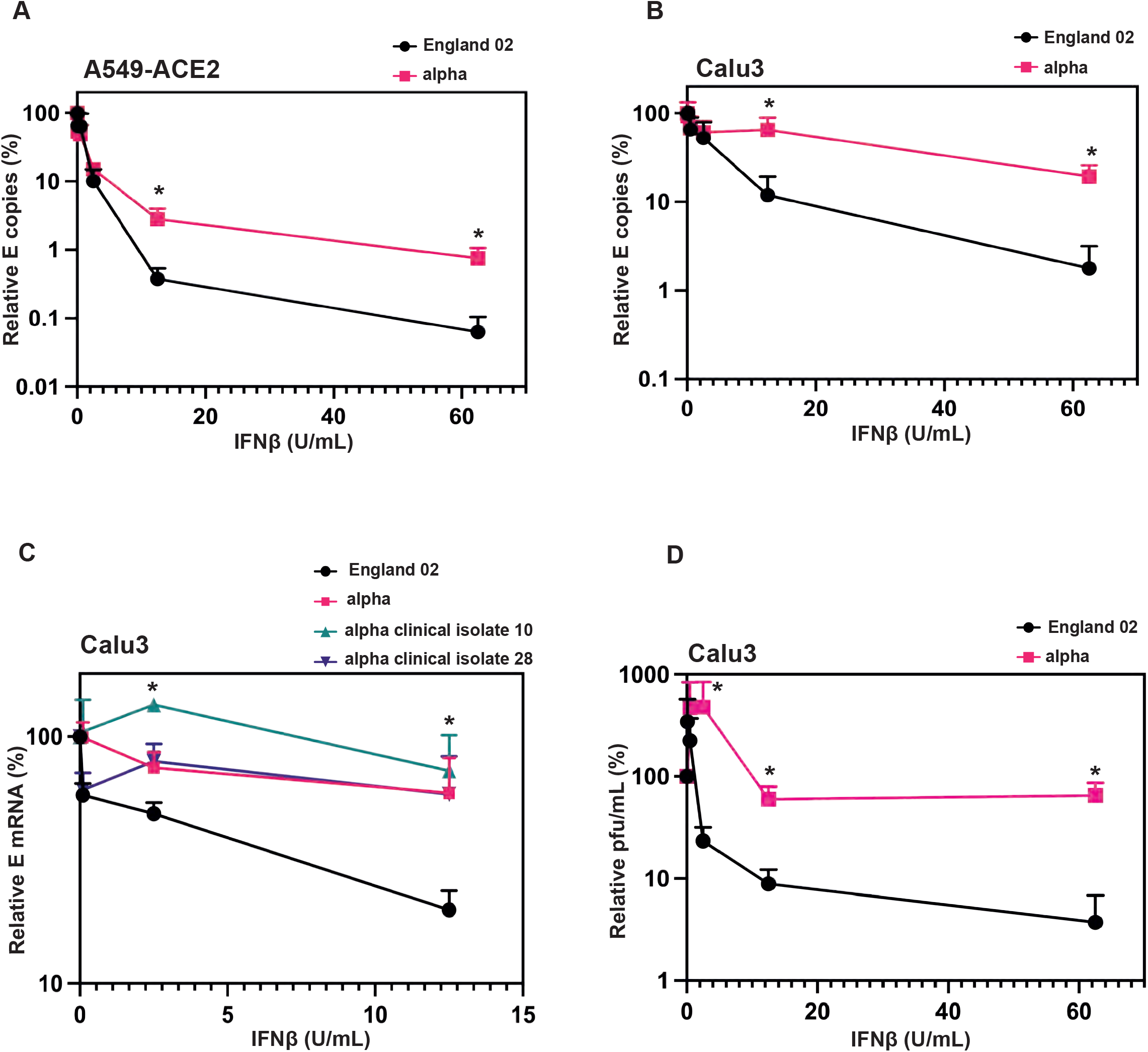
The alpha variant is resistant to IFNβ. A) England 02 and alpha virus infection in A549-ACE2 cells pre-treated with IFNβ. Cells were pre-treated with increasing concentrations of IFNβ for 18 hours prior to infection with either virus at 500 E mRNA copies/cell. Infection was quantified by RT-qPCR of E mRNA from the supernatant 48 hours later and normalised to the un-treated control. B) England 02 and alpha virus infection in Calu-3 cells pre-treated with IFNβ. Cells were pre-treated with increasing concentrations of IFNβ for 18 hours prior infection with either virus at 5000 E copies/cell. Infection was quantified by RT-qPCR of E mRNA from the supernatant 48 hours later and normalised to the un-treated control. C) England 02 and clinical isolates of alpha virus infection in Calu-3 cells pre-treated with IFNβ and harvested as in A and B. Cells were pre-treated with increasing concentrations of IFNβ for 18 hours prior to infection with either virus at 5000 E copies/cell. Infection was quantified by RT-qPCR of cellular E mRNA relative to GAPDH 48 hours later and normalised to the un-treated control. D) Calu-3 cells were infected with England-02 or alpha as in B and supernatant from infected cells used to infect Vero-E6-TMPRSS2 cells for 72 hours. Pfu/ml was determined by plaque assay. Data shown are mean ± SEM, n=3. Statistics were calculated in Prism using *t*-test, stars indicate significance between the different viruses at individual IFN concentrations (*P=<0.05).

### Discordance between the incorporation of furin-processed S proteins into lentiviral particles and native virions

It has been postulated that the P681R and P681H mutations that have emerged in the delta, alpha, and omicron variants enhance S processing, which facilitates a more cell-surface route of entry [33]. However, whether the P681R or P681H mutations confer a greater degree of S processing has been debated [20, 34]. We had previously linked cleavage at the S1/S2 boundary in the Wuhan-1 virus as a factor in reduced IFITM2 sensitivity, and therefore postulated that the P681H mutation may lead to increased S1/S2 cleavage and explain why alpha is IFITM resistant in A549-ACE2s. PLVs assemble at the plasma membrane [35] and incorporate SARS-CoV-2 S into virions that leaches to the cell surface because it escapes COPI-mediated ER/Golgi retention [29], and this process is enhanced by removal of the C-terminal 19 aa of S [36]. By contrast, native CoV virions assemble at, and bud into, intracellular Golgi-derived membranes, and are then secreted. While most studies have compared the incorporation of furin-cleaved S in PLVs versus S in the lysate of SARS-CoV-2 infected cells, we compared S cleavage and incorporation into sucrose-pelleted virions for sequence-verified isolates of the major VOCs, and lentiviral pseudotypes made with the same S (Figure 6A-6F). In contrast to the HEK293T cells producing PLVs, Vero-E6-TMPRSS2 cells infected with fixed doses of the Wuhan-1-like England-02, D614G, alpha, delta and omicron isolates displayed marked difference in cleaved S content in both cells and pelleted virions. Lysates of 30h infected Vero-E6-TMPRSS2 cells displayed markedly higher amounts of the S2 cleavage product as a proportion of uncleaved S for D614G and the VOCs as compared to the Eng-02 isolate. While incorporation of S into harvested virions (S levels in pelleted virions relative to N) was equivalent – Fig 6A and 6C)), virions produced from Vero-E6-TMPRSS2 reflected the cell lysate well: with alpha and omicron showing much higher relative cleaved S incorporation than delta or D614G, which in turn was more pronounced than Eng-02 (Figure 6A-6B). That this contrasts with data from other groups producing virus in other systems highlights that the relative proportion of cleaved S on SARS-CoV-2 virions is likely to be highly dependent on the cell line in which the virus is grown. By contrast, PLVs displayed clear differences with the native virus – while all spikes were similarly expressed in the cell lysates, there were clear differences in the level of PLV incorporation of between PLVs (6D-6F), indicating that PLVs may not give a true reflection of the S conformation on native virions. Discrepancies between lentiviral vectors and virus S processing has also been recently suggested by the Cote group, and it is likely the cell type that the viruses and PLVs are produced in influences the observed S processing and may explain some of the differences in the literature [22, 37]. Given that the structural proteins E, M and N are known to regulate S retention, assembly, and glycosylation [38], we suggest that differences in S cleavage based solely on assays using spike-only PLVs should be interpreted with caution. Furthermore, as demonstrated in Figure 3, PLVs with a ΔCT result in both differential S1/S2 cleavage and cathepsin-dependence, further confirming this needs to be taken into account when determining consequences for spike cleavage.

**Figure 6.**
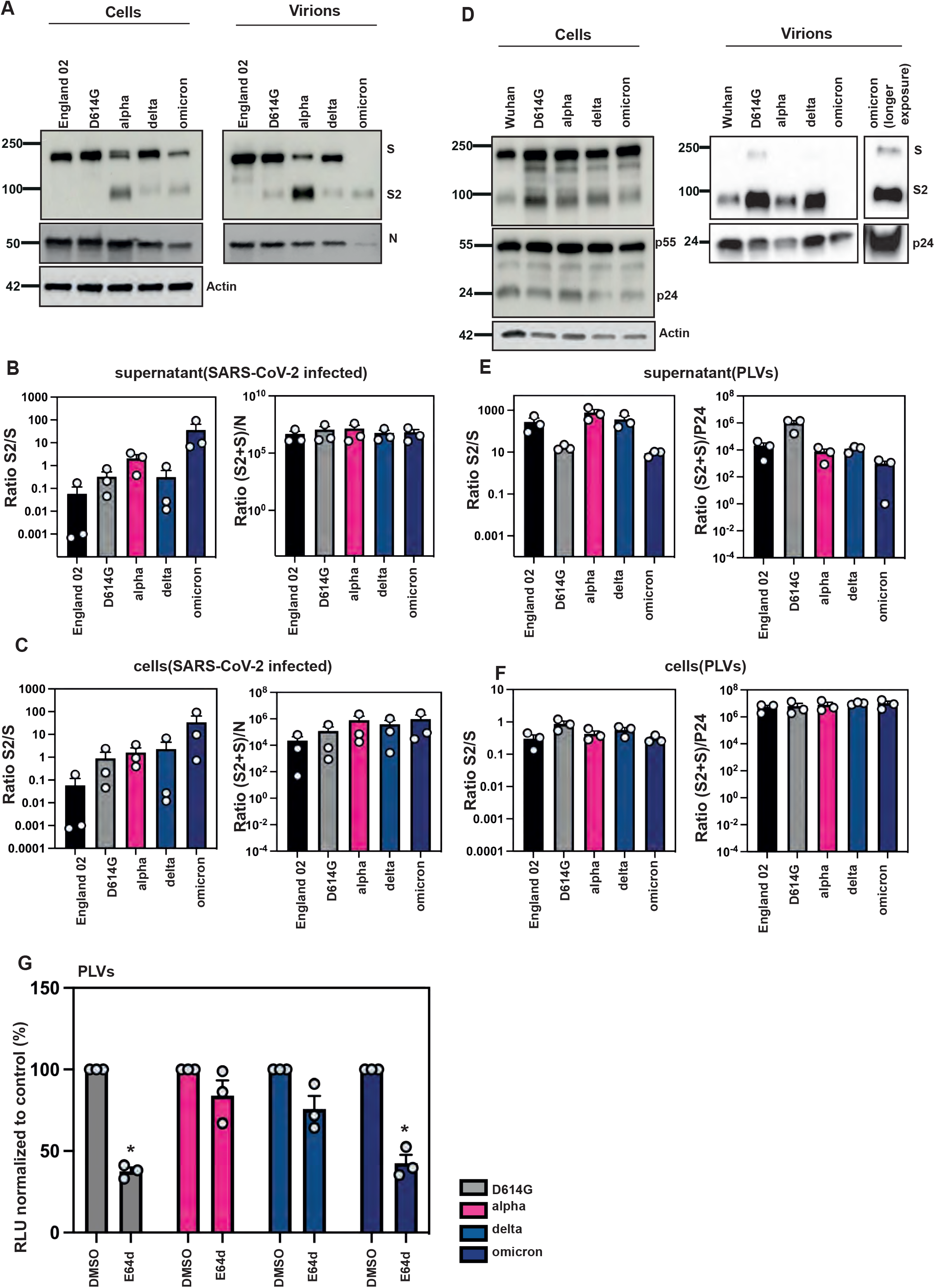
Spike is differentially cleaved across the major variants but not in PLVs. A) representative western blot of spike protein in cell lysates and purified supernatants of Vero-E6-TMPRSS2 infected cells. Cells were infected with Wuhan, D614G, alpha, delta or omicron isolates at an MOI of 1 for 30h. Virions were purified through a 20% sucrose gradient. B) representative western blot of spike protein in cell lysates and purified supernatants from PLVs. PLVs were produced in 293T/17s and immunoblotted 48h after transfection. Virions were purified through a 20% sucrose gradient. C) Quantification of spike in cell lysates of infected Vero-E6-TMPRSS2 cells after 30h. Graph shows ratio of S2 over total S, and ratio of S2 and total S over N. D) Quantification of spike in purified supernatant from infected Vero-E6-TMPRSS2 cells after 30h. Graph shows ratio of S2 over total S, and ratio of S2 and total S over N. E) Quantification of spike in cell lysates of 293T/17 cells used to produce PLVs. Graph shows ratio of S2 over total S, and ratio of S2 and total S over p55. F) Quantification of spike in purified PLVs produced in 293T/17 cells. Graph shows ratio of S2 over total S, and ratio of S2 and total S over p24. G) E64d sensitivity of D614G, alpha, delta and omicron PLVs. A549-ACE2 were pre-treated with 10*μ*M E64d for 1 hour prior to transduction and infection quantified by Luciferase activity 48h later. Data shown are mean ± SEM, n=3. Statistics were calculated in Prism using *t*-test, stars indicate significance between control and drug (*P=<0.05).

Next, we tested if the alpha, delta and omicron variants used the same route of entry given the polybasic cleavage site mutations. Other groups have suggested that the omicron variant, despite containing a P681H mutation, is more dependent on the endosomal route of entry [27] which may account for why omicron retains IFITM sensitivity. We hypothesised that despite the P681H mutation, the myriad of mutations in the RBD that confer a more “closed” conformation forces it towards a cathepsin-dependent route of entry. To test this, we pre-treated A549-ACE2 cells with the endosomal cathepsin inhibitor E64d and infected with PLVs of D614G, alpha, delta and omicron. We found that, in line with what others have described, omicron displayed similar E64d sensitivity to D614G (Figure 6G). The alpha or delta variants essentially showed no significant dependence on cathepsin-mediated S cleavage relative to D614G. Overall, these results suggest S1/S2 cleavage is highly cell-type dependent and does not necessarily correlate with route of viral entry.

### The P681H mutation is necessary for conferring IFITM and IFNβ resistance in alpha by promoting a near cell surface route of viral entry

Our previous data indicated IFITM sensitivity of SARS-CoV-2 S can be increased by deletion of the polybasic cleavage site [14]. Given that the alpha S acquired the P681H mutation and we demonstrated that it is relatively insensitive to an inhibitor of endosomal entry (Figure 6G), we hypothesized that P681H might be a determinant of resistance to IFN and IFITMs for the alpha S. Using PLVs on A549-ACE2-IFITM cells, we first confirmed that ablation of the entire polybasic cleavage site increases IFITM2 sensitivity to D614G, as we have previously demonstrated for the Wuhan-1 S [14]. As expected, D614GΔPRRA is highly sensitive to IFITM2 and is not cleaved on PLV particles (Figure 7A, S3C, S3D, S3E). Next, we tested if the same polybasic cleavage site deletion sensitized alpha to the IFITMs. Not only was the ΔHRRA mutant sensitive to IFITM2, we also abolished the IFITM3 enhancement phenotype observed with the alpha PLV (Figure 7A, further statistics in S3A, S3B) suggesting that the furin cleavage site was essential for both of these phenotypes. Having confirmed that alpha S could be sensitized to IFITM2 by deletion of the HRRA site, we next tested whether the P681H mutation alone could confer IFITM resistance to a D614G S, and vice versa. We found that the P681H mutation in the D614G background was sufficient to abolish IFITM2 sensitivity, however was not able to confer the same level of IFITM3-mediated enhancement we observe with alpha. However, the H681P mutation in alpha sensitized the alpha PLV to IFITM2, although not to the same extent as the ΔHRRA mutation, and also reduced the IFITM3 enhancement of alpha. We noted that the H681P mutation did not revert the cleavage of the alpha S in the context of PLVs (Figure S3C, S3D, S3E), however as suggested in Figure 5, making conclusions on S cleavage from PLVs may not represent the real virus. We concluded that although the P681H mutation is necessary for IFITM resistance, it is likely that other contextual mutations in the alpha S are required for this to be sufficient for IFITM3 enhancement. Next, we tested whether the P681R mutation in the D614G background alters IFITM sensitivity (S). Unlike the P681H mutation, the P681R mutation did not alter the IFITM sensitivity of D614G (Figure S3F). Additionally, reverting the R861 to a P in the delta S had little impact on IFITM sensitivity (Figure S3G), further suggesting that the P681R mutation cannot confer IFITM resistance.

**Figure 7.**
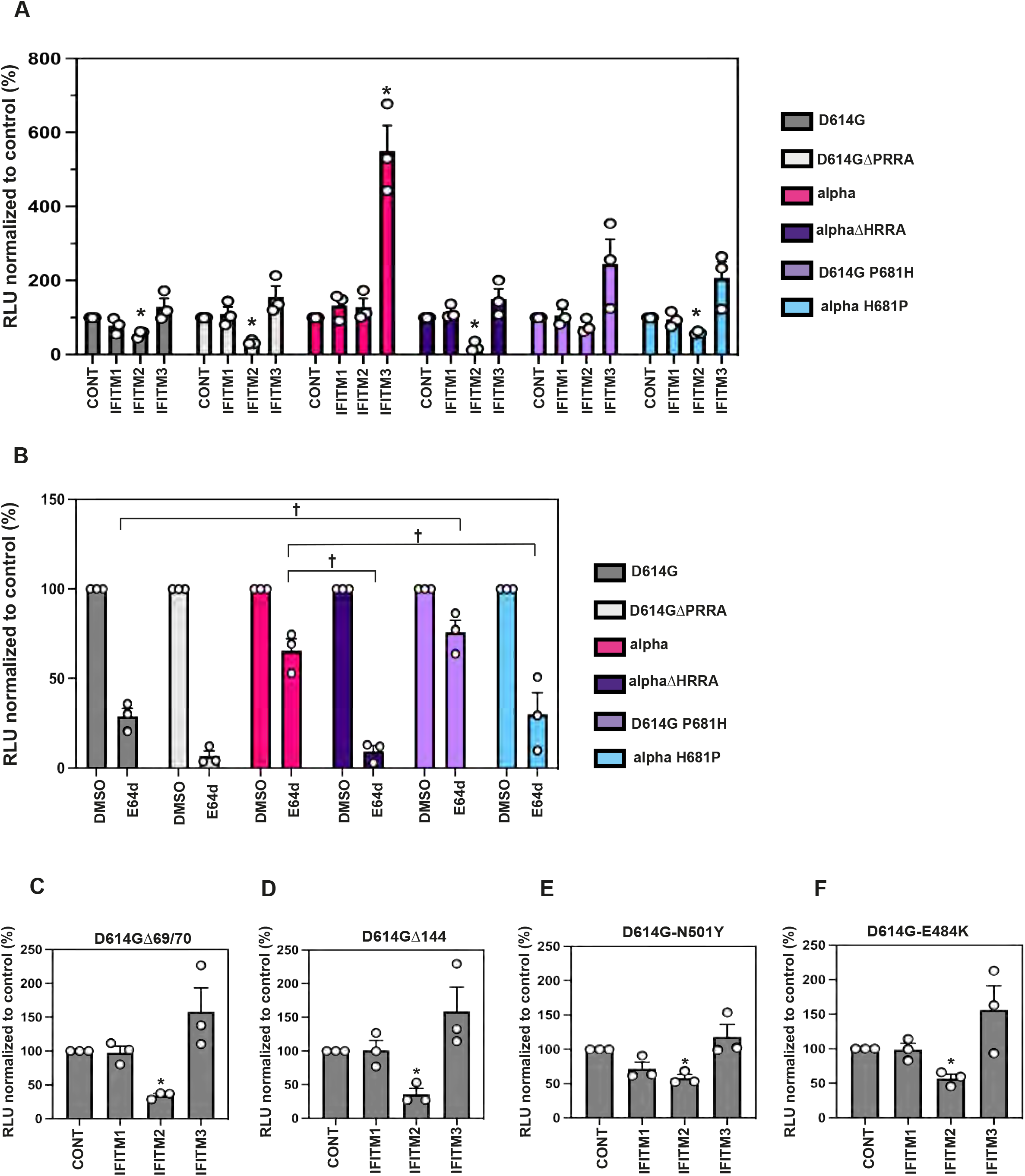
The P681H mutation confers IFITM resistance to a wild-type spike and a reduced dependence on E64d. A) D614G, D614GΔPRRA, alpha, alpha-ΔHRRA, D614G P681H, and alpha H681P and PLVs infection in A549-ACE2 cells stably expressing the individual IFITMs. PLVs entry was quantified by Luciferase activity 48 hours later and normalized to control cells. B) E64d treatment of A549-ACE2 cells infected with PLVs. A549-ACE2s were pre-treated with 10*μ*M E64d prior to transduction with D614G, alpha, delta or omicron PLVs and infection quantified by Luciferase activity 48h later. C, D, E, F) PLVs of individual mutations from alpha in the D614G background were used to infect A549-ACE2-IFITM cells and infection quantified by Luciferase activity 48 hours later. Infection normalised to control cells with no IFITM is shown. Data shown are mean ± SEM, n=3. Statistics were calculated in Prism using *ANOVA*, stars indicate significance between control cell and individual IFITM or drug and crosses indicate statistical significance between different IFITM/drug conditions (*P=<0.05).

We then wanted to confirm whether the P681H mutation confers IFITM resistance by reducing the preference for endosomal entry in the A549-ACE2 system. Previously, we demonstrated that the alpha S is relatively insensitive to the effects of the cathepsin inhibitor E64d. To test whether the increased IFITM sensitivity of the ΔHRRA and H681P mutants correlated with increased endosomal entry and therefore exposure to IFITM2, we pre-treated cells with the cathepsin inhibitor E64d as before, and infected with PLVs (Figure 7D). As expected, we found that the D614G and alpha polybasic cleavage site deletions were highly sensitive to E64d. Additionally, the H681P mutation conferred increased E64d sensitivity to alpha, suggesting this mutant is more reliant on cathepsin-dependent entry and therefore more likely to encounter IFITM2. As expected, the inverse P681H mutation in the context of D614G conferred decreased E64d sensitivity to the wild-type S. This suggests that the P681H mutation alone is sufficient to confer increased preference for cell-surface entry to a D614G S. In the context of the alpha S, we further suggest that the P681H mutation is a determinant of route of viral entry and therefore sensitivity to antiviral proteins that occupy endosomal compartments. Having established that the P681R mutation did not alter IFITM sensitivity, we hypothesized that this mutation alone would not reduce E64d sensitivity to a wild-type S. Indeed, while the P681H mutation reduces cathepsin-dependence, the P681R mutation is indistinguishable from D614G in terms of E64d sensitivity (Figure S3H). This suggests that the P681R mutation does not confer cell-surface mediated entry in the A549-ACE2 cells. Finally, to confirm if any of the other defining mutations in the alpha spike altered IFITM sensitivity, we generated single mutants of the Δ69/70 (Figure 7C), Δ144 (7D), N501Y (7E) and the E484K mutation (7F), which emerged in several sub-lineages. None of these mutations significantly altered IFITM resistance.

### Reversion of the P681H mutation sensitises the alpha variant to IFNβ and IFITM2

Finally, we tested if the H681P reversion was sufficient to revert the overall IFNβ resistance phenotype of alpha. We constructed a recombinant molecular clone of SARS-CoV-2 Wuhan-1 encoding S from the alpha variant. This virus essentially mimicked the resistance of the alpha variant itself to IFNβ in comparison to England-02, demonstrating that the alpha S alone is sufficient to confer a level of IFN-I resistance in A549-ACE2 cells (Figure 8A). Then, we reverted from this recombinant virus the amino acid residue H681 to a proline. Importantly, this single point mutation was sufficient to confer a significant sensitivity to IFNβ in Calu3 cells indicating it was a major determinant of IFN resistance in alpha S (Figure 8B). Furthermore, we wanted to confirm whether siRNA knockdown of IFITM2 was sufficient to rescue the IFNβ sensitivity of the Wuhan(B.1.1.7 Spike H681P) virus. We confirmed that IFITM2 knockdown had no effect on other ISG expression and IFNβ signalling, measured by STAT1 phosphorylation and Viperin expression (Figure 8C). We showed that the H681P reverted virus was rescued from IFNβ restriction by IFITM2 knockdown, meanwhile the Wuhan(B.1.1.7 Spike) virus was unaffected, consistent with this virus being resistant to IFITM restriction (Figure 8D). Thus, this confirmed that the S protein of the alpha variant of SARS-CoV-2 is a determinant of type-I IFN resistance, which is primarily modulated by IFITM2. Most importantly, the P681H mutation is necessary for this. Interestingly, when we immunoblotted purified virions of the Wuhan(B.1.1.7 Spike) and Wuhan(B.1.1.7 Spike-H681P), we found, similar to the PLVs (Figure 6B-C), that the H681P reversion did not affect the cleavage of the alpha S (Figure 8E). Thus, P681H mutation is a major determinant of IFN Type I resistance in the alpha variant but it must exert its activity downstream of cleavage itself.

**Figure 8.**
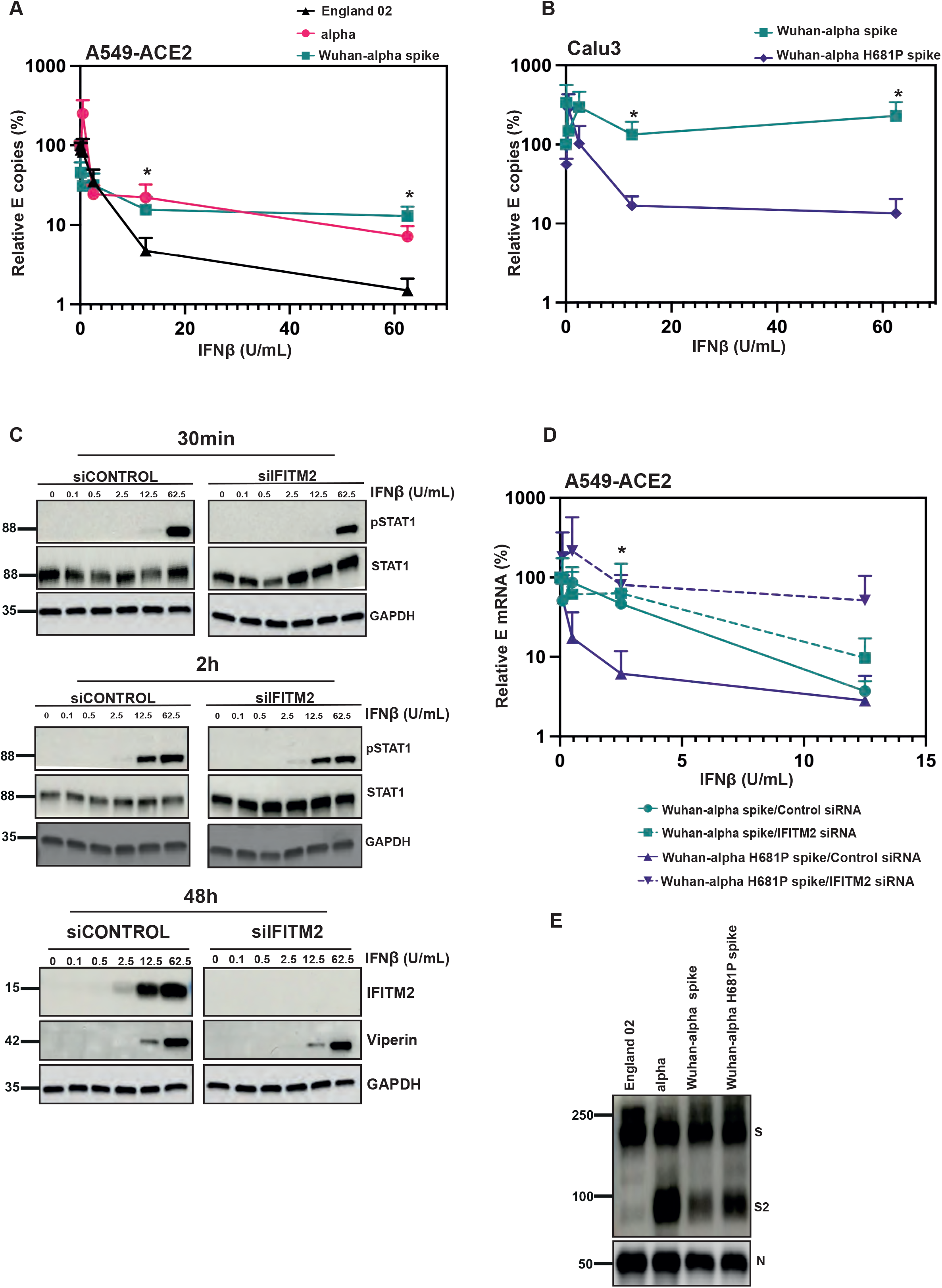
The P681H mutation is necessary and sufficient for IFNβ resistance. A) England 02, alpha, and Wuhan(alpha spike) virus infection in A549-ACE2 cells pretreated with IFNβ. Cells were pre-treated with increasing concentrations of IFNβ for 18 hours prior to infection with either virus at 500 E copies/cell. Infection was quantified by RT-qPCR of E mRNA in the supernatant 48 hours later and normalised to the mock control. B) Wuhan(alpha spike) and Wuhan(alpha spike H681P) virus infection in Calu-3 cells pre-treated with IFNβ. Cells were pre-treated with increasing concentrations of IFNβ for 18 hours prior to infection with either virus at 5000 E copies/cell. Infection was quantified by RT-qPCR of E mRNA in the supernatant 48 hours later and normalised to the mock control. C) representative immunoblot of pSTAT1 and STAT1 in cell lysates knocked down for IFITM2 or a non-targeting control and subsequently treated with IFNβ. A549-ACE2 cells were transfected with siRNAs against non-targeting control or IFITM2 and then treated with IFNβ for 30 minutes or 2 hours and immunoblotted for pSTAT1 and STAT1, or treated for 24 hours and blotted for viperin and IFITM2. D) A549-ACE2 cells were transfected with siRNAs against non-targeting control or IFITM2 for 24 hours and then treated with IFNβ for 18 hours prior to infection with Wuhan(alpha spike) or Wuhan(alpha spike H681P) at 500 copies/cell. Infection was quantified by RT-qPCR of E gene relative to GAPDH 48 hours later; graph represents E mRNA levels relative to GAPDH. Data shown are mean ± SEM, n=3. Statistics were calculated in Prism using t-test, black stars indicate significance at each IFNβ concentration between the different viruses (*P=<0.05). E) Representative immunoblot of England-02, alpha, Wuhan(alpha spike) and Wuhan(alpha spike H681P viral stocks. England-02, alpha, Wuhan(alpha spike), Wuhan(alpha spike H681P) viruses were purified through 20% sucrose and immunoblotted for spike and N proteins.

## DISCUSSION

Here we have shown that the S protein of the alpha variant of SARS-CoV-2 is a determinant of viral resistance to IFN-I. This maps to the histidine residue adjacent to the polybasic cleavage site that has been mutated from the parental proline. While this has been shown to enhance S cleavage at the S1/S2 boundary in a context dependent manner [39], the H at this position in alpha rather than the cleavage itself appears to confer the IFN-resistance phenotype. This is reinforced by the finding that deleting the last 19 amino acids of D614G spike results in enhanced S1/S2 cleavage, but not enhanced IFITM resistance, further suggesting that cleavage per se is not the determining factor of the alpha variants IFITM resistance. The P681H mutation correlates with the abolition of the residual sensitivity to endosomal cathepsin inhibitors implying a change in viral entry route that distinguishes alpha from delta. This residue is also necessary to confer both resistance to IFITM2 and enhancement by IFITM3, and as we demonstrated in our previous study [14], confirms that the polybasic cleavage site can modulate IFITM entry restriction. Furthermore, we demonstrate that this mutation alone in a wild-type D614G S is sufficient to promote reduced IFITM sensitivity, whilst the delta P681R mutation is not. Furthermore, we note that infection by alpha is enhanced in the presence of IFITM3, and this is abolished by cyclosporin H, cytoplasmic tail deletion, or the H681P mutation. IFITM3 has previously been reported to enhance the entry of the coronavirus OC43, and more recently has been suggested to enhance the entry of Hepatitis B and D [24, 40]. Although it is surprising that an antiviral protein can enhance infection, this phenotype in multiple viruses suggests a common mechanism of hijacking host factors for viral entry. We also see a degree of IFITM enhancement by 1 or 2 with the alpha virus and PLVs to a variable degree. We suggest that this may be a factor of the trafficking of IFITMs through multiple compartments and the occasional presence of IFITMs 1 and 2 in the compartment IFITM3 usually resides. Enhancement of coronavirus entry by the mutant IFITM1Δ117-125 has previously been documented, suggesting that IFITM1 can enhance viral entry depending on localisation [41]. We have also previously shown that the Y19F mutation in IFITM2 also results in enhancement of Wuhan entry, further suggesting that IFITM localisation can alter their capacity to enhance coronavirus infection [14].

We suggest that the P681H change in alpha changes the site of viral fusion, therefore avoiding the endosomal compartment where IFITM2 predominantly resides. Consistent with this, we showed that the alpha S in a PLV is less sensitive to the cathepsin inhibitor E64d. Thus, we propose that these changes in the alpha S have, in part, arisen to resist innate immunity. At least two studies suggest that variants of SARS-CoV-2 have begun to evolve further resistance to interferon-induced innate immunity [31, 42, 43]. In one, viral isolates over the pandemic showed a reduced sensitivity to type I interferons in culture [42]; in a second the alpha variant has a significantly reduced propensity to trigger pattern recognition in epithelial cells by cytoplasmic RNA sensors [31, 43]. In contrast, another study shows no difference in IFN sensitivity of the new variants in African green monkey Vero-E6 cells [44], although species-specificity in viral sensitivity to ISGs is a well characterized trait that could explain this discrepancy [45]. The SARS-CoV-2 genome contains multiple mechanisms to counteract host innate immune responses, and much remains to be learned about the mechanisms deployed by this virus and its relatives. While many reports on SARS-CoV-2 evolution have naturally focussed on the pressing concern of potential for vaccine escape, it is very unlikely that all selective adaptations that we see arising in VOCs can be solely due to escape from adaptive immunity. The alpha S, for example, only displays a minor reduction in sensitivity to neutralizing antibodies (NAbs) [8, 46-48]. However, this VOC had a considerable transmission advantage, with suspicions that it may have arisen in an immunocompromised individual with a persistent infection giving ample time for changes to be selected that further evade innate immunity, including those that target viral entry [49, 50].

In terms of IFITM resistance of VOC S proteins, so far we have only seen marked change in phenotype for the alpha variant. This is despite the fact that both delta and omicron, variants that superseded alpha, also showed an adaptation for enhanced S1/S2 cleavage with a P681R and P681H change respectively [20, 39]. This would suggest that cleavage of S1/S2 is necessary but not sufficient for IFITM resistance, and that other mutations in each cognate S act in concert to determine relative IFITM sensitivity. Despite the increased cleavage of a D614G-containing isolate, delta and omicron relative to England-02, these viruses are not IFITM resistant. This suggests the P681H mutation confers IFITM resistance through a mechanism distinct from S1/S2 cleavage itself. We and others show that omicron is sensitive to E64d inhibition, and we suggest this preference for endosomal entry correlates with omicron’s IFITM sensitivity [27]. The S of omicron contains 30 mutations, 12 of which are in the RBD and have been suggested to increase the affinity for ACE2 [51]. The constellation of mutations in the RBD of omicron also promotes an “RBD down” closed conformation, which necessitates cathepsin-mediated cleavage in the endosome rather than surface TMPRSS2-mediated cleavage [27]. This suggests that the up-conformation of the RBD is required for H681 to exert its IFN-resistance phenotype [27]. Furthermore, omicron contains a H655Y mutation which has been suggested to enhance endosomal entry [52]. Despite delta containing a P681R mutation, we report that this S is not IFITM resistant, nor is the kappa variant which also bears a P681R mutation and was relatively short-lived as a variant. Despite the delta S demonstrating E64d insensitivity, the P681R mutation alone does not result in reduced IFITM sensitivity or decreased E64d sensitivity to a D614G S. This implies that there are other factors besides the P681R mutation governing delta’s route of viral entry. Two recent papers have suggested that certain matrix metalloproteinases (MMPs) can mediate an alternative route of entry to TMPRSS2, and that this can be utilised by the delta variant [22, 53]. It is possible that a culmination of these viral entry routes are variably present in different cell types, and may therefore explain differential IFITM sensitivities by VOCs. Finally, the delta variant also contains different RBD mutations than alpha, in particular the T478K and L452R mutations, which may also affect the RBD conformation and be a factor in delta’s relative sensitivity to IFITMs. The mutations in the RBD’s of delta and omicron have led to hypotheses that both of these variants were driven by antibody escape, suggesting selection pressures on the alpha variant may have been more due to innate immunity. It is important to note that the discordance between virion incorporated S species in the native SARS-CoV-2 particle and lentiviral pseudotypes imply a degree of cell type dependency as well as cellular location of viral assembly in the relative presence of cleaved S. We also demonstrate that this is something of particular concern for those using C-terminal deletions of the COP-I retention signal in S. We would be cautious describing some of the phenotypes associated with VOC S protein simply to differences in furin-cleavage efficiency, or phenotypes ascribed from Δ19aa PLVs when the route of viral entry is implicit to the phenotype.

Viral glycoproteins are dynamic structures that shift through large-scale conformational changes while interacting with their cognate receptors mediating viral membrane fusion [54]. Such context dependency is therefore likely to be complex and will arise under competing selective pressures. Indeed, we have previously shown that the HIV-1 envelope glycoprotein of transmitted viruses is IFITM insensitive and this contributes to their overall type I IFN resistance [55]. As HIV-1 infection progresses over the first 6 months in an infected person, the circulating variants increase in IFN/IFITM sensitivity and this is determined by adaptive changes in Env that resist the early neutralizing antibody response [56]. Such escape has structural and functional implications for such dynamic proteins that may impact upon receptor interactions and route of entry into the target cell.

The mapping of IFN-I resistance to P681H to the polybasic cleavage site of alpha, but yet reversion of the IFN-I sensitivity by the restoration of the P without affecting the cleavage of virion associated S, suggests that the H681 is exerting its effects on viral entry and IFITM/IFN-I sensitivity downstream of cleavage itself. While it is possible that this could be simply related to stability of the cleaved form, it is intriguing to note that the C-terminal RRAR of S1 has also been proposed as a ligand for neuropilin-1 (NRP-1), a receptor for furin-processed growth factors like VEGF-A. NRP-1 was found to promote the entry and replication of SARS-CoV-2 in an FCS-dependent manner [57, 58]. Given the accumulating evidence that inter-protomer interactions in the S trimer affect the accessibility of cleavage sites in S [59], future studies will determine whether a role for NRP-1 in entry is also governing sensitivity to IFITM-restriction and IFN-sensitivity.

While the furin cleavage site of the SARS-CoV-2 S reduces its IFITM sensitivity, other interferon-induced proteins may contribute to this phenotype. The guanylate binding protein family, and particularly GBP2 and GBP5, have been shown to have a general antiviral activity against enveloped viruses by dysregulating furin processing of diverse viral and cellular proteins [60]. Similarly, IFITM overexpression in HIV-infected cells can lead to their incorporation into virions and in some cases promote defects in glycoprotein incorporation [61]. Future studies will confirm whether either of these mechanisms are involved in the IFN-resistance associated with the P681H mutation in alpha [27].

In summary, the S protein of SARS-CoV-2 alpha increases resistance to IFN-I and this correlates with the P681H mutation. Furthermore, this correlates with IFITM resistance as IFITM2 knockdown rescues the IFN sensitive alpha H681P virus, but not alpha. Despite also containing P681R/P681H mutations, the delta and omicron variants are not IFITM resistant in the A549-ACE2 system. We suggest that factors such as RBD conformation and alternate routes of viral entry all act in concert to determine the relative sensitivities of S proteins to antiviral proteins that affect viral entry.

## MATERIALS AND METHODS

### Cells and plasmids

HEK293T-17 (ATCC, CRL-11268^™^), Calu-3 (ATCC, HTB-55^™^), A549-ACE2, Vero-E6, Vero-E6-TMPRSS2 and A549-ACE2 expressing the individual IFITM proteins were cultured in DMEM (Gibco) with 10% FBS (Invitrogen) and 200μg/ml Gentamicin (Sigma), and incubated at 37°C, 5% CO2. ACE2, TMPRSS2, and IFITM stable overexpression cells were generated as previously described [14].

Codon optimised SARS-CoV-2 Wuhan Spike and ACE2 were kindly given by Dr. Nigel Temperton. Codon optimised variant Spikes (alpha, beta) were kindly given by Dr. Katie Doores. Codon optimised variant Spikes (gamma, kappa, delta) were kindly given by Professor Wendy Barclay. Plasmid containing TMPRSS2 gene was kindly given by Dr. Caroline Goujon. Spike mutants were generated with Q5® Site-Directed Mutagenesis Kit (E0554) following the manufacturer’s instructions, and using the following forward and reverse primers: D614G (GCTGTACCAGGGCGTGAATTGCA, ACGGCCACCTGATTGCTG) B.1.351. Δ242-244 (ATTTCATATCTTACACCAGGC, ATGCAGGGTCTGGAATCTG) D614G P681H (GACCAATAGCcacAGAAGAGCCAGAAGC, TGGGTCTGGTAGCTGGCG) B117 ΔHRRA (AGAAGCGTGGCCAGCCAG, GCTATTGGTCTGGGTCTGGTAG) B117 H681P (GACCAATAGCcccAGAAGAGCCAG, TGGGTCTGGTAGCTGGCG) Δ 69/70 (AGCGGCACCAATGGCACC, GATGGCGTGGAACCAGGTC), D144 (CATAAGAACAACAAGAGC, ATAAACACCCAGGAAAGG) A549 stable cell lines expressing ACE2 (pMIGR1-puro), and IFITMs (pLHCX) were generated and selected as described previously [14].

### Production of Pseudotyped Lentiviral Vectors (PLVs) and infection

HEK293T-17 cells were transfected with firefly luciferase expressing vector (CSXW), HIV gag-pol (8.91) and Spike plasmid with PEI-max as previously described [14]. 100ul of viral supernatant was then used to transduce each cell line of interest and readout measured by Luciferase activity 48 hours later (Promega Steady-Glo® (E2550)).

### Cyclosporin H assay

Cells were pre-treated with 30µM of Cyclosporin H (Sigma, SML1575) for 18 hours. Cells were then infected with PLVs as above and viral entry quantified by Luciferase activity 48 hours later.

### E64d assay

A549-ACE2 cells were pre-treated with 10µM of E64d (Sigma; E8640) for 1h at 37°C prior to infection. Cells were transduced with PLVs and infection determined by luciferase activity 48 hours later.

### Passage and titration of SARS-CoV-2

PHE England strain 02/2020 and D614G isolate were propagated in Vero-E6-TMPRSS2 cells and titre was determined by plaque assay [14]. Plaque assays were performed by infecting Vero-E6-TMPRSS2 with serial dilutions of SARS-CoV-2 for 1h. Subsequently, 2X overlay media (DMEM + 2% FBS + 0.1% agarose) was added, and infected cells were fixed with 4% PFA 72 hours after infection and stained with Crystal Violet. Plaques were counted and multiplicity of infection calculated for subsequent experiments. A replication-competent alpha variant was kindly provided by Professor Wendy Barclay (Imperial College London)[62]. All virus stocks were sequence confirmed in the Spike gene at each passage to ensure no loss of the FCS.

### Generation of recombinant full-length viruses

We used the previously described Transformation-Associated Recombination (TAR) in yeast method [63], with some modifications, to generate the mutant viruses described in this study. Briefly, a set of overlapping cDNA fragments representing the entire genomes of SARS-CoV-2 Wuhan isolate (GenBank: MN908947.3) and the B.1.1.7 alpha variant were chemically synthesized and cloned into pUC57-Kan (Bio Basic Canada Inc and Genewiz, respectively). The cDNA fragment representing the 5’ terminus of the viral genome contained the bacteriophage T7 RNA polymerase promoter preceded by a short sequence stretch homologous to the *Xho*I-cut end of the TAR in yeast vector pEB2 [64]. The fragment representing the 3’ terminus contained the T7 RNA polymerase termination sequences followed by a short segment homologous to the *Bam*HI-cut end of pEB2.

To generate Wuhan virus carrying the alpha variant spike, a mixture of the relevant synthetic cDNA fragments of the Wuhan and alpha variants was cotransformed with *Xho*I-*BamHI*-cut pEB2 into the *Saccharomyces cerevisiae* strain TYC1 (MATa, ura3-52, leu2Δ1, cyh2^r^, containing a knockout of DNA Ligase 4) [64] that had been made competent for DNA uptake using the LiCl2-based Yeast transformation kit (YEAST1-1KT, Merck). The transformed cells were plated on minimal synthetic defined (SD) agar medium lacking uracil (Ura) but containing 0.002% (w/v) cycloheximide to prevent selection of cells carrying the empty vector. Following incubation at 30°C for 4 to 5 days, colonies of the yeast transformants were screened by PCR using specific primers to identify those carrying plasmid with fully assembled genomes. Selected positive colonies were then expanded to grow in 200 ml SD-Ura dropout medium and the plasmid extracted. Approximately 4 µg of the extracted material was then used as template to *in vitro* synthesized viral genomic RNA transcripts using the Ribomax T7 RNA transcription Kit (Promega) and Ribo m7G Cap Analogue (Promega) as per the manufacturer’s protocol. Approximately 2.5 µg of the *in vitro* synthesized RNA was used to transfect ∼6 ×10^5^ BHK-hACE2-N cells stably expressing the SARS-CoV-2 N and the human ACE2 genes [65] using the MessengerMax lipofection kit (Thermo Scientific) as per the manufacturer’s instructions. Cells were then incubated until signs of viral replication (syncytia formation) became visible (usually after 2-3 days), at which time the medium was collected (P0 stock) and used further as a source of rescued virus to infect Vero-E6 cells to generate P1 and P2 stocks. Full genome sequences of viruses collected from from P0 and P1 stocks were obtained in order to confirm the presence of the desired mutations and exclude the presence of other spurious mutations. Viruses were sequenced using Oxford Nanopore as previously described [66].

To generate Wuhan virus carrying alpha spike gene with the H681P mutation, we first introduced this mutation into the relevant alpha variant cDNA fragment by site-directed mutagenesis. This fragment was combined with those described above and the mixture was then used to generate plasmid pEB2 carrying the cDNA genome of Wuhan encoding the alpha spike H681P by the TAR in yeast procedure. The virus rescue and subsequent characterisation were performed as described above.

### Isolation and Propagation of Clinical Viral Isolates

Viruses were isolated on Vero-E6 cells (ATCC CRL 1586™) from combined naso-oropharyngeal swabs submitted for routine diagnostic testing by real-time RT-PCR and shown to be from the B.1.1.7 (alpha) variant by on-site whole-genome sequencing (Oxford Nanopore Technologies, Oxford, UK) [67]. Infected cells were cultured at 37°C and 5% CO2, in Dulbecco’s modified Eagle’s medium (DMEM, Gibco^™^, Thermo Fisher, UK) supplemented with 2% foetal bovine serum (FBS, Merck, Germany), pen/strep and amphotericin B.

All work performed with full-length SARS-CoV-2 preparations, as well as isolation and propagation of viral isolates from swabs, was conducted inside a class II microbiological safety cabinet in a biosafety level 3 (BSL3) facility at King’s College London.

### Infection with replication competent SARS-CoV-2

1.5×10^5^ A549-ACE2 cells were infected for 1 hour at 37°C with SARS-CoV-2 replication competent viruses at MOI 0.01 or 500 E gene mRNA copies/cell. 2×10^5^ Calu-3 cells were infected for 1h at 37°C with SARS-CoV-2 replication competent viruses at 5000 E gene mRNA copies/cell. Media was replaced and cells were incubated for 48 hours at 37°C, after which cells or supernatant were harvested for RNA extraction or protein analysis.

### Intracellular N staining

1.5×10^5^ A549-ACE2 IFITM cells were infected for 1 hour at 37°C with SARS-CoV-2 replication competent VOCs to achieve the same percentage of infected cells in the mock condition. After 24h infection cells were trypsinized and fixed with 4% PFA during 30 min at RT. Cells were permeabilized with 1X PBS + 0.5 % Triton during 10 min following blocking with 5% FBS in 1X PBS during 20 min. After blocking cells were stained with anti-N antibody (CR3009, mouse) during 45 min at RT and washed once with 1X PBS. Next, cells were incubated with secondary anti-mouse alexa488 antibody for 25 min. Finally, cells were washed with 1X PBS and analyzed on a BD FACS Canto II using FloJo software.

### Interferon assays

Cells were treated with different doses of IFNβ (PBL Assay Science, 11415-1) for 18 hours prior to infection. The following day media was replaced, and the infection performed as described above. Viral RNA levels in cells or supernatants were measured 48 hours after infection by RT-qPCR.

### siRNA knockdown of IFITM2

A549-ACE2 cells were reverse transfected using 20pmol of Non-targeting siRNA (D-001206-13-20) or IFITM2 siRNA (M-020103-02-0010) with 1*μ*L of RNAi max (Invitrogen). Cells were incubated for 24h prior to a second round of reverse transfection. 8h later, cells were treated with different doses of IFNβ. Following 18h of IFN treatment cells were infected with full-length viruses as previously described.

### RT-qPCR

RNA from infected cells was extracted using QIAGEN RNeasy (QIAGEN RNeasy Mini Kit, 74106) following the manufacturer’s instructions. 1*μ*L of each extracted RNA was used to performed one step RT-qPCR using TaqMan Fast Virus 1-Step Master Mix (Invitrogen). The relative quantities of envelope (E) gene were measured using SARS-CoV-2 (2019-nCoV) CDC qPCR Probe Assay (IDT DNA technologies). Relative quantities of E gene were normalized to GAPDH mRNA levels (Applied Bioscience, Hs99999905_m1).

Supernatant RNA was extracted using RNAdvance Viral XP (Beckman) following the manufacturer’s instructions. 5*μ*L of each RNA was used for one-step RT-qPCR (TaqMan™ Fast Virus 1-Step Master Mix) to measured relative quantities of E and calibrated to a standard curve of E kindly provided by Professor Wendy Barclay.

### SDS-PAGE and Western blotting

Cellular samples were lyzed in reducing Laemmli buffer at 95°C for 10 minutes. Supernatant or viral stock samples were centrifuged at 18,000 RCF through a 20% sucrose cushion for 1 hour at 4°C prior to lysis in reducing Laemmli buffer. Samples were separated on 8–16 % Mini-PROTEAN® TGX™ Precast gels (Bio-Rad) and transferred onto nitrocellulose membrane. Membranes were blocked in milk or BSA prior to detection with specific antibodies: 1:1000 ACE2 rabbit (Abcam, Ab108209), 1:5000 GAPDH rabbit (Abcam, Ab9485), 1:2000 anti-GAPDH mouse (Proteintech, 60004-1-Ig), 1:5000 HSP90 mouse (Genetex, Gt×109753), 1:50 HIV-1 p24Gag mouse (48 ref before) 1:1000 Spike mouse (Genetex, Gtx632604), 1:1000 anti-SARS-CoV-2 N rabbit (GeneTex, GTX135357) 1:1000 anti-pSTAT1 mouse (BD Transduction Laboratories, 612133), 1:1000 anti-STAT1 rabbit (Cell Signalling, 9172S), 1:1000 antiviperin mouse (Millipore, MABF106). Proteins were detected using LI-COR and ImageQuant LAS 4000 cameras.

### Ethics

Clinical samples were retrieved by the direct care team in the Directorate of Infection, at St Thomas Hospital, London, UK, and anonymized before sending to the King’s College London laboratories for virus isolation and propagation. Sample collection and studies were performed in accordance with the UK Policy Framework for Health and Social Care Research and with specific Research Ethics Committee approval (REC 20/SC/0310).

## ACKNOWLEDGEMENTS

We are grateful to Nigel Temperton, Caroline Goujon, Katie Doores, Wendy Barclay and Public Health England for reagents. We acknowledge the G2P-UK National Virology consortium funded by MRC/UKRI (grant ref: MR/W005611/1) and the Barclay Lab at Imperial College London for providing the alpha variant. We thank E. J. Louis, University of Leicester for generously providing the TAR in yeast system. Finally, we thank all other members of the Neil and Swanson groups for their helpful advice and provision of sugar-based support.

## AUTHORS CONTRIBUTION

Experiments were performed by MJL, HW, AD and HDW. SP, RPG, LS and GN collected, sequenced and isolated clinical viral isolates. MP, AHP, GDL, VMC, WF, NS, and RO generated reverse genetics-derived viruses. MJL, HW, AD and HDW analysed data. CMS provided reagents, funding support and advice. HW and SJDN wrote the manuscript. All authors edited the manuscript and provided comments.

## FIGURE LEGENDS

**Supplementary Figure 1.**
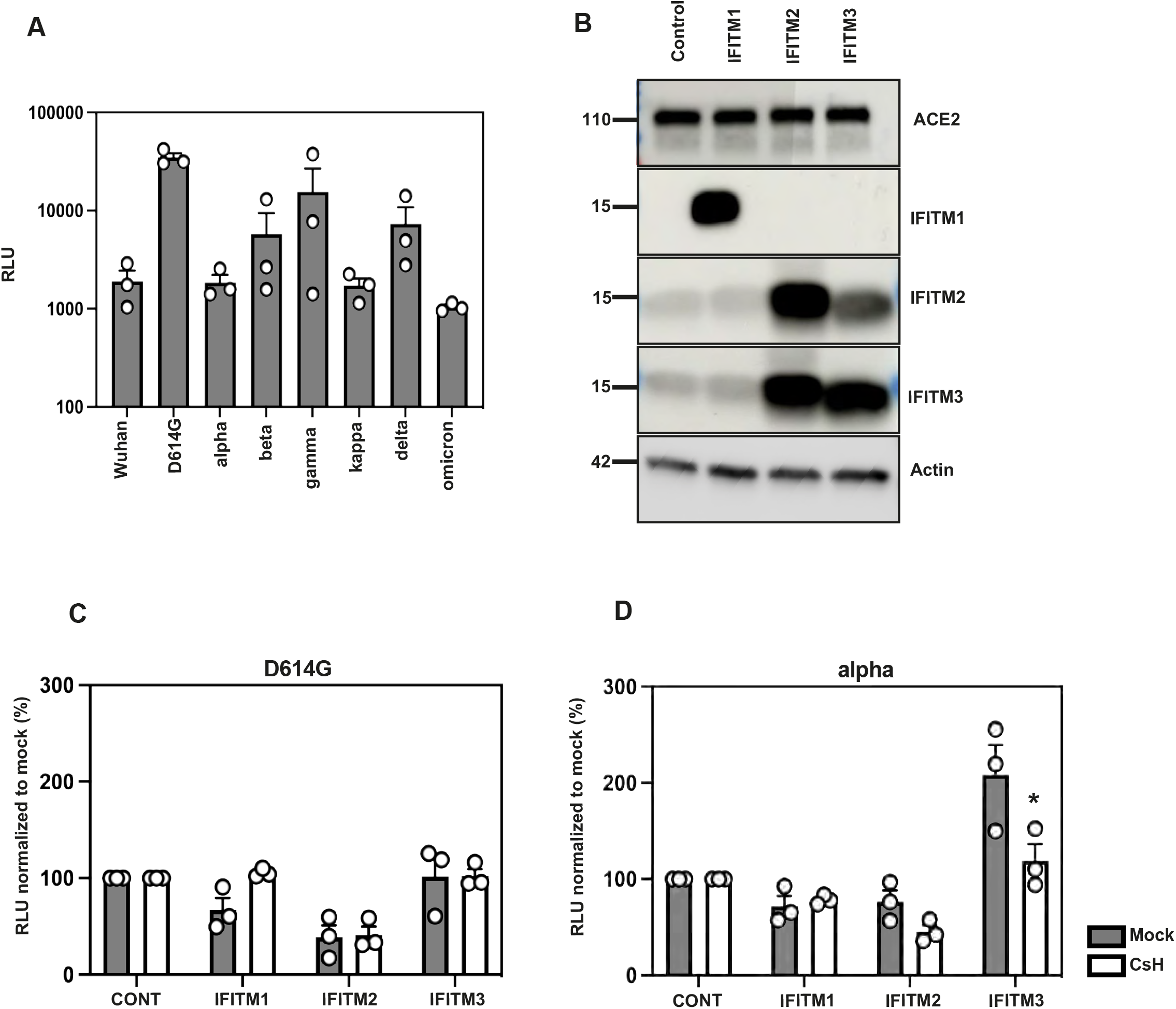
Cyclosporin H treatment abolishes IFITM3 enhancement. A) Relative titre of VOC PLVs on A549-ACE2 cells. A549-ACE2 were transduced with PLVs of Wuhan, D614G, alpha, beta, gamma, kappa, delta and omicron for 48h and infectivity quantified by Luciferase activity. B) Representative immunoblot of A549-ACE2 cells stably expressing IFITMs 1, 2 and 3. **C, D)** D614G PLVs pre-treated with Cyclosporin H. A549-ACE2s stably expressing the individual IFITMs were pre-treated with 30 µM of Cyclosporin H for 18 hours prior to infection with D614G PLVs. Infection was quantified by Luciferase activity 48 hours after infection and normalized to control cells. Data shown are mean ± SEM, n=3. Statistics were calculated in Prism using *t*-test, stars indicate significance between IFITM3 mock and IFITM3 CsH (*P=<0.05).

**Supplementary Figure 2.**
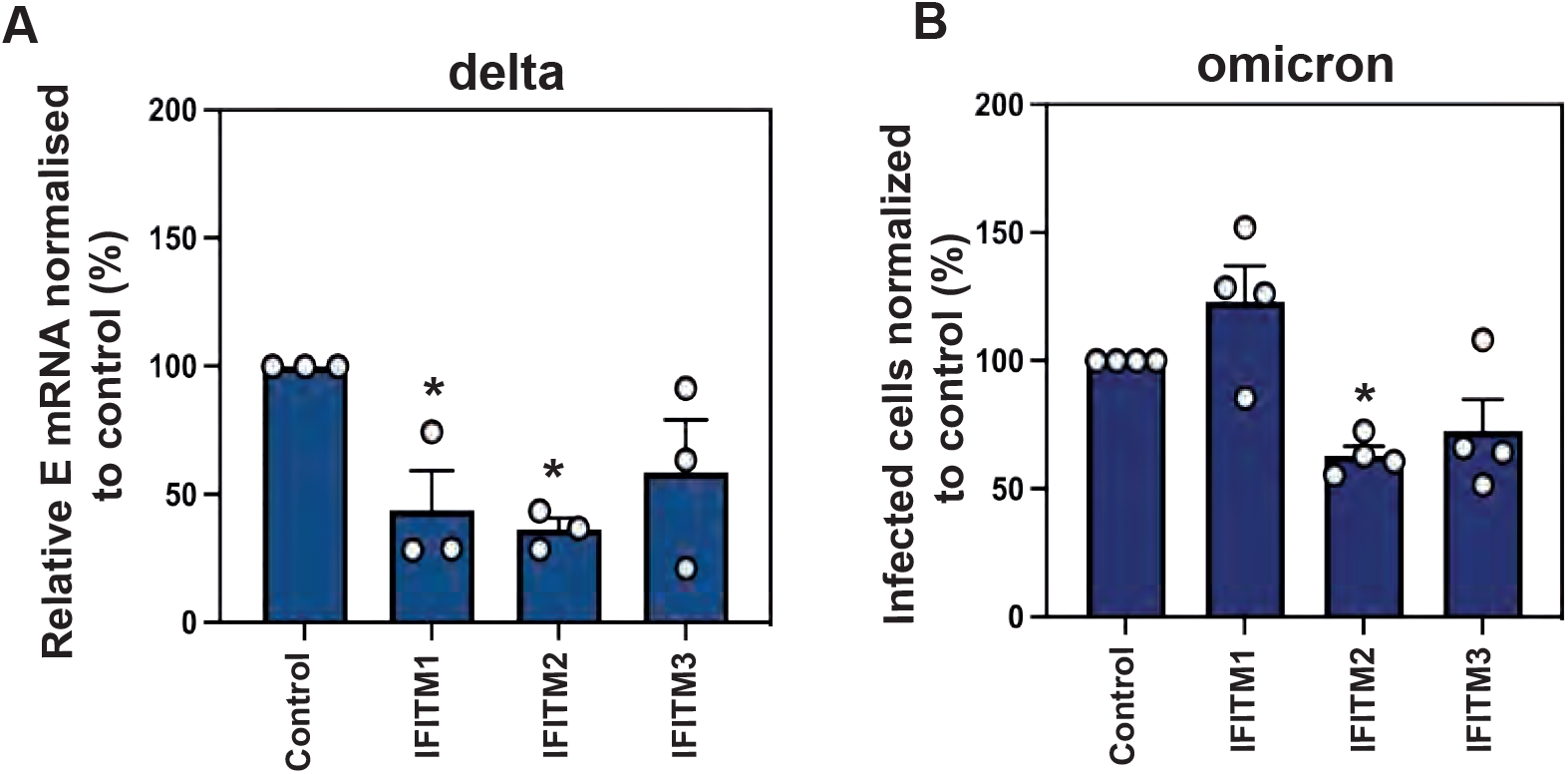
The delta and omicron viruses are IFITM sensitive in A549-ACE2s. A) Infection of A549-ACE2 stably expressing the individual IFITMs with delta virus at MOI 0.01. Infection was quantified by RT-qPCR of E gene relative to GAPDH 48 hours later; graph represents E mRNA levels relative to GAPDH. B) intracellular N staining by flow cytometry of A549-ACE2 IFITM cells infected with omicron virus. A549-ACE2 expressing the individual IFITMs were infected with an omicron isolate for 48h. Infection was measured by percentage of N positive cells by flow cytometry. Data analysed in FlowJo. Data shown are mean ± SEM, n=3. Statistics were calculated in Prism using *t*-test, stars indicate significance between control cell and individual IFITM (*P=<0.05).

**Supplementary Figure 3.**
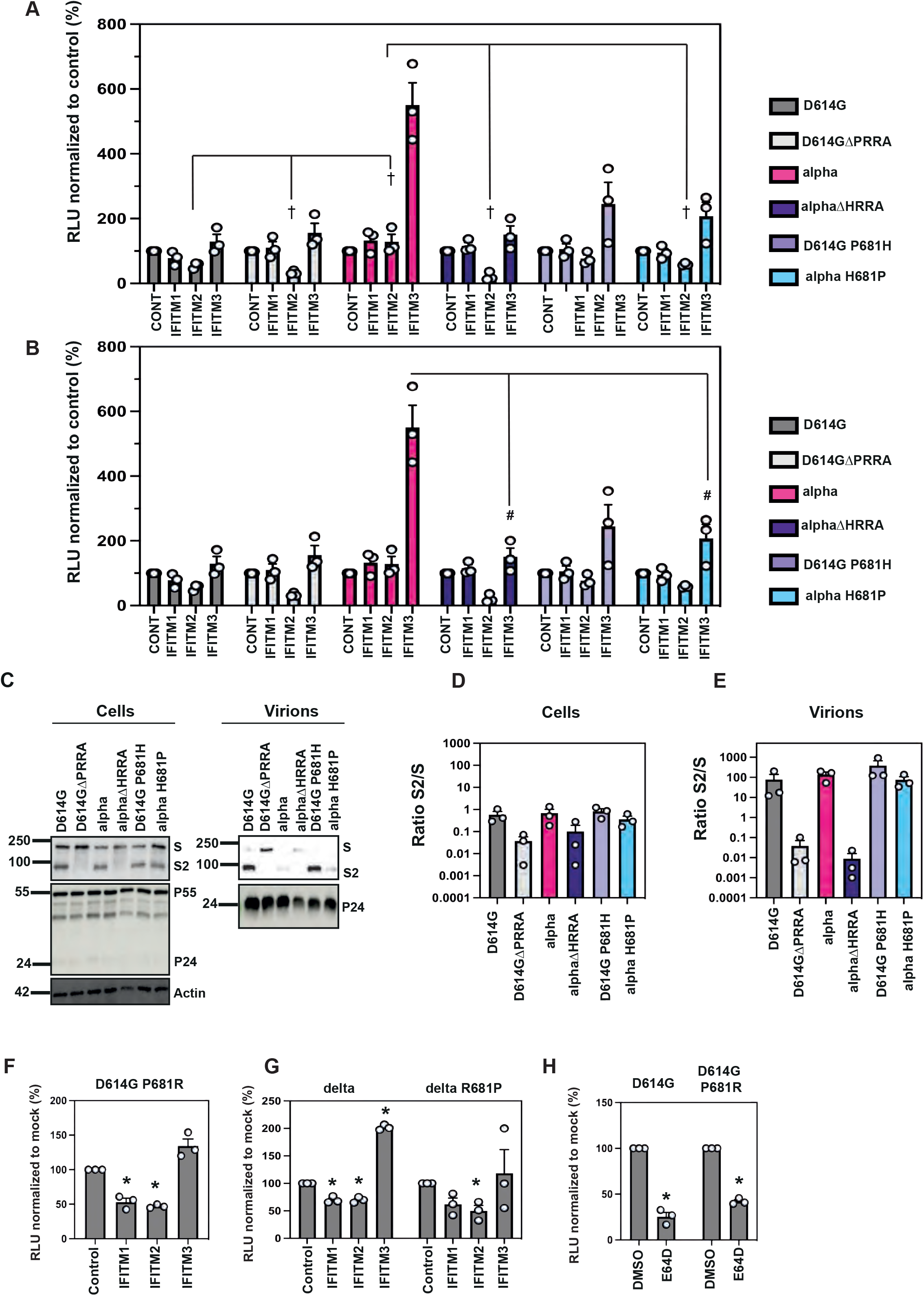
The P681R mutation does not confer IFITM resistance. A) cross symbol shows statistical significance by t-test between IFITM2 of D614G mutants compared to D614G, and statistical significance by t-test between IFITM2 of alpha mutants compared to the alpha of Figure 6. B) Hashtag symbol shows statistical significance by t-test between IFITM3 of the alpha mutants compared to the alpha of Figure 6. C) Representative immunoblots of cell lysates and purified supernatants of PLV production in 293T/17 cells. Virions were purified through a 20% sucrose cushion. D, E) Quantification of S2 over total S of cell lysates (D) and supernatant of PLVs produced in 293T/17 cells (E). F) A549-ACE2 cells stably expressing the individual IFITMs were infected with D614G P681R PLVs. PLV entry was quantified by Luciferase activity 48 hours after infection and infectivity normalized to control cells. **G)** A549-ACE2 cells stably expressing the individual IFITMs were infected with delta or delta R681P PLVs. PLV entry was quantified by Luciferase activity 48 hours after infection and infectivity normalized to control cells. **H)**. A549-ACE2s were pre-treated with 10*μ*M E64d prior to transduction with D614G or D614G P681R PLVs and infection quantified by Luciferase activity 48h later. Data shown are mean ± SEM, n=3. Statistics were calculated in Prism using *t*-test, stars indicate significance between mock and individual IFITM or drug (*P=<0.05).

## REFERENCES

1. Hoffmann M, Kleine-Weber H, Pohlmann S. A Multibasic Cleavage Site in the Spike Protein of SARS-CoV-2 Is Essential for Infection of Human Lung Cells. Mol Cell. 2020;78(4):779-84 e5. Epub 2020/05/05. doi: 10.1016/j.molcel.2020.04.022. PubMed PMID: 32362314; PubMed Central PMCID: PMCPMC7194065.

2. Peacock TP, Goldhill DH, Zhou J, Baillon L, Frise R, Swann OC, et al. The furin cleavage site in the SARS-CoV-2 spike protein is required for transmission in ferrets. Nat Microbiol. 2021;6(7):899-909. Epub 2021/04/29. doi: 10.1038/s41564-021-00908-w. PubMed PMID: 33907312.

3. Lindstrom JC, Engebretsen S, Kristoffersen AB, Ro GOI, Palomares AD, Engo-Monsen K, et al. Increased transmissibility of the alpha SARS-CoV-2 variant: evidence from contact tracing data in Oslo, January to February 2021. Infect Dis (Lond). 2021:1-6. Epub 2021/10/08. doi: 10.1080/23744235.2021.1977382. PubMed PMID: 34618665.

4. Tanaka H, Hirayama A, Nagai H, Shirai C, Takahashi Y, Shinomiya H, et al. Increased Transmissibility of the SARS-CoV-2 Alpha Variant in a Japanese Population. Int J Environ Res Public Health. 2021;18(15). Epub 2021/08/08. doi: 10.3390/ijerph18157752. PubMed PMID: 34360046; PubMed Central PMCID: PMCPMC8345780.

5. Mok BW, Liu H, Deng S, Liu J, Zhang AJ, Lau SY, et al. Low dose inocula of SARS-CoV-2 Alpha variant transmits more efficiently than earlier variants in hamsters. Commun Biol. 2021;4(1):1102. Epub 2021/09/22. doi: 10.1038/s42003-021-02640-x. PubMed PMID: 34545191; PubMed Central PMCID: PMCPMC8452646.

6. Meng B, Kemp SA, Papa G, Datir R, Ferreira I, Marelli S, et al. Recurrent emergence of SARS-CoV-2 spike deletion H69/V70 and its role in the Alpha variant B.1.1.7. Cell Rep. 2021;35(13):109292. Epub 2021/06/25. doi: 10.1016/j.celrep.2021.109292. PubMed PMID: 34166617; PubMed Central PMCID: PMCPMC8185188.

7. Chi X, Yan R, Zhang J, Zhang G, Zhang Y, Hao M, et al. A neutralizing human antibody binds to the N-terminal domain of the Spike protein of SARS-CoV-2. Science. 2020;369(6504):650-5. Epub 20200622. doi: 10.1126/science.abc6952. PubMed PMID: 32571838; PubMed Central PMCID: PMCPMC7319273.

8. Graham C, Seow J, Huettner I, Khan H, Kouphou N, Acors S, et al. Neutralization potency of monoclonal antibodies recognizing dominant and subdominant epitopes on SARS-CoV-2 Spike is impacted by the B.1.1.7 variant. Immunity. 2021;54(6):1276-89 e6. Epub 2021/04/10. doi: 10.1016/j.immuni.2021.03.023. PubMed PMID: 33836142; PubMed Central PMCID: PMCPMC8015430.

9. Mohammad A, Abubaker J, Al-Mulla F. Structural modelling of SARS-CoV-2 alpha variant (B.1.1.7) suggests enhanced furin binding and infectivity. Virus Res. 2021;303:198522. Epub 2021/07/28. doi: 10.1016/j.virusres.2021.198522. PubMed PMID: 34314772; PubMed Central PMCID: PMCPMC8310422.

10. Zhang L, Mann M, Syed Z, Reynolds HM, Tian E, Samara NL, et al. Furin cleavage of the SARS-CoV-2 spike is modulated by O-glycosylation. bioRxiv. 2021. Epub 2021/02/11. doi: 10.1101/2021.02.05.429982. PubMed PMID: 33564758; PubMed Central PMCID: PMCPMC7872346.

11. Rajah MM, Hubert M, Bishop E, Saunders N, Robinot R, Grzelak L, et al. SARS-CoV-2 Alpha, Beta, and Delta variants display enhanced Spike-mediated syncytia formation. EMBO J. 2021;40(24):e108944. Epub 20211025. doi: 10.15252/embj.2021108944. PubMed PMID: 34601723; PubMed Central PMCID: PMCPMC8646911.

12. Araf Y, Akter F, Tang YD, Fatemi R, Parvez MSA, Zheng C, et al. Omicron variant of SARS-CoV-2: Genomics, transmissibility, and responses to current COVID-19 vaccines. J Med Virol. 2022;94(5):1825-32. Epub 20220123. doi: 10.1002/jmv.27588. PubMed PMID: 35023191; PubMed Central PMCID: PMCPMC9015557.

13. Ostrov DA, Knox GW. Emerging mutation patterns in SARS-CoV-2 variants. Biochem Biophys Res Commun. 2022;586:87-92. Epub 20211122. doi: 10.1016/j.bbrc.2021.11.059. PubMed PMID: 34837837; PubMed Central PMCID: PMCPMC8606318.

14. Winstone H, Lista MJ, Reid AC, Bouton C, Pickering S, Galao RP, et al. The Polybasic Cleavage Site in SARS-CoV-2 Spike Modulates Viral Sensitivity to Type I Interferon and IFITM2. J Virol. 2021;95(9). Epub 2021/02/11. doi: 10.1128/JVI.02422-20. PubMed PMID: 33563656; PubMed Central PMCID: PMCPMC8104117.

15. Shi G, Kenney AD, Kudryashova E, Zani A, Zhang L, Lai KK, et al. Opposing activities of IFITM proteins in SARS-CoV-2 infection. EMBO J. 2021;40(3):e106501. Epub 2020/12/04. doi: 10.15252/embj.2020106501. PubMed PMID: 33270927; PubMed Central PMCID: PMCPMC7744865.

16. Bailey CC, Zhong G, Huang IC, Farzan M. IFITM-Family Proteins: The Cell’s First Line of Antiviral Defense. Annu Rev Virol. 2014;1:261-83. Epub 2015/01/20. doi: 10.1146/annurev-virology-031413-085537. PubMed PMID: 25599080; PubMed Central PMCID: PMCPMC4295558.

17. Shi G, Schwartz O, Compton AA. More than meets the I: the diverse antiviral and cellular functions of interferon-induced transmembrane proteins. Retrovirology. 2017;14(1):53. Epub 2017/11/23. doi: 10.1186/s12977-017-0377-y. PubMed PMID: 29162141; PubMed Central PMCID: PMCPMC5697417.

18. Jia R, Pan Q, Ding S, Rong L, Liu SL, Geng Y, et al. The N-terminal region of IFITM3 modulates its antiviral activity by regulating IFITM3 cellular localization. J Virol. 2012;86(24):13697-707. Epub 20121010. doi: 10.1128/JVI.01828-12. PubMed PMID: 23055554; PubMed Central PMCID: PMCPMC3503121.

19. Jia R, Xu F, Qian J, Yao Y, Miao C, Zheng YM, et al. Identification of an endocytic signal essential for the antiviral action of IFITM3. Cell Microbiol. 2014;16(7):1080-93. Epub 2014/02/14. doi: 10.1111/cmi.12262. PubMed PMID: 24521078; PubMed Central PMCID: PMCPMC4065222.

20. Liu Y, Liu J, Johnson BA, Xia H, Ku Z, Schindewolf C, et al. Delta spike P681R mutation enhances SARS-CoV-2 fitness over Alpha variant. bioRxiv. 2021. Epub 2021/09/01. doi: 10.1101/2021.08.12.456173. PubMed PMID: 34462752; PubMed Central PMCID: PMCPMC8404900.

21. Liu Y, Liu J, Johnson BA, Xia H, Ku Z, Schindewolf C, et al. Delta spike P681R mutation enhances SARS-CoV-2 fitness over Alpha variant. Cell Rep. 2022;39(7):110829. Epub 20220429. doi: 10.1016/j.celrep.2022.110829. PubMed PMID: 35550680; PubMed Central PMCID: PMCPMC9050581.

22. Mehdi Benlarbi GL, Corby Fink, Kathy Fu, Rory P. Mulloy, Alexandra Phan, Ardeshir Ariana, Corina M. Stewart, Jérémie Prévost, Guillaume Beaudoin-Bussières, Redaet Daniel, Yuxia Bo, Julien Yockell-Lelièvre, William L. Stanford, Patrick M. Giguère, Samira Mubareka, Andrés Finzi, Gregory A. Dekaban, Jimmy D. Dikeakos, Marceline Côté. Identification of a SARS-CoV-2 host metalloproteinase-dependent entry pathway differentially used by SARS-CoV-2 and variants of concern Alpha, Delta, and Omicron. bioRxiv. 2022.

23. Korber B, Fischer WM, Gnanakaran S, Yoon H, Theiler J, Abfalterer W, et al. Tracking Changes in SARS-CoV-2 Spike: Evidence that D614G Increases Infectivity of the COVID-19 Virus. Cell. 2020;182(4):812-27 e19. Epub 20200703. doi: 10.1016/j.cell.2020.06.043. PubMed PMID: 32697968; PubMed Central PMCID: PMCPMC7332439.

24. Zhao X, Guo F, Liu F, Cuconati A, Chang J, Block TM, et al. Interferon induction of IFITM proteins promotes infection by human coronavirus OC43. Proc Natl Acad Sci U S A. 2014;111(18):6756-61. Epub 2014/04/23. doi: 10.1073/pnas.1320856111. PubMed PMID: 24753610; PubMed Central PMCID: PMCPMC4020042.

25. Prelli Bozzo C, Nchioua R, Volcic M, Koepke L, Kruger J, Schutz D, et al. IFITM proteins promote SARS-CoV-2 infection and are targets for virus inhibition in vitro. Nat Commun. 2021;12(1):4584. Epub 2021/07/30. doi: 10.1038/s41467-021-24817-y. PubMed PMID: 34321474; PubMed Central PMCID: PMCPMC8319209.

26. Petrillo C, Thorne LG, Unali G, Schiroli G, Giordano AMS, Piras F, et al. Cyclosporine H Overcomes Innate Immune Restrictions to Improve Lentiviral Transduction and Gene Editing In Human Hematopoietic Stem Cells. Cell Stem Cell. 2018;23(6):820-32 e9. Epub 20181108. doi: 10.1016/j.stem.2018.10.008. PubMed PMID: 30416070; PubMed Central PMCID: PMCPMC6292841.

27. Dejan Mesner A-KR, Matthew V.X Whelan, Taylor Bronzovich, Tafhima Haider, Lucy G. Thorne, Greg J. Towers, Clare Jolly. SARS-CoV-2 Spike evolution influences GBP and IFITM sensitivity. bioRxiv. 2022.

28. Chen HY, Huang C, Tian L, Huang X, Zhang C, Llewellyn GN, et al. Cytoplasmic Tail Truncation of SARS-CoV-2 Spike Protein Enhances Titer of Pseudotyped Vectors but Masks the Effect of the D614G Mutation. J Virol. 2021;95(22):e0096621. Epub 20210908. doi: 10.1128/JVI.00966-21. PubMed PMID: 34495700; PubMed Central PMCID: PMCPMC8549521.

29. Jackson CB, Farzan M, Chen B, Choe H. Mechanisms of SARS-CoV-2 entry into cells. Nat Rev Mol Cell Biol. 2022;23(1):3-20. Epub 20211005. doi: 10.1038/s41580-021-00418-x. PubMed PMID: 34611326; PubMed Central PMCID: PMCPMC8491763.

30. Jouvenet NG, C.; Banerjee, A. Clash of the titans: interferons and SARS-CoV-2. Trends in Immunology. 2021.

31. Thorne LG, Bouhaddou M, Reuschl AK, Zuliani-Alvarez L, Polacco B, Pelin A, et al. Evolution of enhanced innate immune evasion by SARS-CoV-2. Nature. 2022;602(7897):487-95. Epub 20211223. doi: 10.1038/s41586-021-04352-y. PubMed PMID: 34942634; PubMed Central PMCID: PMCPMC8850198.

32. Guo K, Barrett BS, Mickens KL, Vladar EK, Morrison JH, Hasenkrug KJ, et al. Interferon Resistance of Emerging SARS-CoV-2 Variants. bioRxiv. 2021. Epub 20211210. doi: 10.1101/2021.03.20.436257. PubMed PMID: 33758840; PubMed Central PMCID: PMCPMC7986999.

33. Takeda M. Proteolytic activation of SARS-CoV-2 spike protein. Microbiol Immunol. 2022;66(1):15–23. Epub 20211012. doi: 10.1111/1348-0421.12945. PubMed PMID: 34561887; PubMed Central PMCID: PMCPMC8652499.

34. Lubinski B, Fernandes MHV, Frazier L, Tang T, Daniel S, Diel DG, et al. Functional evaluation of the P681H mutation on the proteolytic activation the SARS-CoV-2 variant B.1.1.7 (Alpha) spike. bioRxiv. 2021. Epub 20211101. doi: 10.1101/2021.04.06.438731. PubMed PMID: 33851153; PubMed Central PMCID: PMCPMC8043443.

35. Ramdas P, Sahu AK, Mishra T, Bhardwaj V, Chande A. From Entry to Egress: Strategic Exploitation of the Cellular Processes by HIV-1. Front Microbiol. 2020;11:559792. Epub 20201204. doi: 10.3389/fmicb.2020.559792. PubMed PMID: 33343516; PubMed Central PMCID: PMCPMC7746852.

36. Roy S, Ghani K, de Campos-Lima PO, Caruso M. A stable platform for the production of virus-like particles pseudotyped with the severe acute respiratory syndrome coronavirus-2 (SARS-CoV-2) spike protein. Virus Res. 2021;295:198305. Epub 20210119. doi: 10.1016/j.virusres.2021.198305. PubMed PMID: 33482242; PubMed Central PMCID: PMCPMC7817443.

37. Kreutzberger AJB, Sanyal A, Saminathan A, Bloyet LM, Stumpf S, Liu Z, et al. SARS-CoV-2 requires acidic pH to infect cells. bioRxiv. 2022. Epub 20220614. doi: 10.1101/2022.06.09.495472. PubMed PMID: 35702155; PubMed Central PMCID: PMCPMC9196115.

38. Xu R, Shi M, Li J, Song P, Li N. Construction of SARS-CoV-2 Virus-Like Particles by Mammalian Expression System. Front Bioeng Biotechnol. 2020;8:862. Epub 20200730. doi: 10.3389/fbioe.2020.00862. PubMed PMID: 32850726; PubMed Central PMCID: PMCPMC7409377.

39. Peacock TP, Sheppard CM, Brown JC, Goonawardane N, Zhou J, Whiteley M, et al. The SARS-CoV-2 variants associated with infections in India, B.1.617, show enhanced spike cleavage by furin. bioRxiv. 2021:2021.05.28.446163. doi: 10.1101/2021.05.28.446163.

40. Palatini M, Muller SF, Kirstgen M, Leiting S, Lehmann F, Soppa L, et al. IFITM3 Interacts with the HBV/HDV Receptor NTCP and Modulates Virus Entry and Infection. Viruses. 2022;14(4). Epub 20220330. doi: 10.3390/v14040727. PubMed PMID: 35458456; PubMed Central PMCID: PMCPMC9027621.

41. Zhao X, Sehgal M, Hou Z, Cheng J, Shu S, Wu S, et al. Identification of Residues Controlling Restriction versus Enhancing Activities of IFITM Proteins on Entry of Human Coronaviruses. J Virol. 2018;92(6). Epub 2017/12/22. doi: 10.1128/JVI.01535-17. PubMed PMID: 29263263; PubMed Central PMCID: PMCPMC5827390.

42. Guo K, Barrett BS, Mickens KL, Hasenkrug KJ, Santiago ML. Interferon Resistance of Emerging SARS-CoV-2 Variants. bioRxiv. 2021. Epub 2021/03/25. doi: 10.1101/2021.03.20.436257. PubMed PMID: 33758840; PubMed Central PMCID: PMCPMC7986999.

43. Thorne LG, Bouhaddou M, Reuschl AK, Zuliani-Alvarez L, Polacco B, Pelin A, et al. Evolution of enhanced innate immune evasion by the SARS-CoV-2 B.1.1.7 UK variant. bioRxiv. 2021. Epub 2021/06/16. doi: 10.1101/2021.06.06.446826. PubMed PMID: 34127972; PubMed Central PMCID: PMCPMC8202424.

44. Michael Rajah M, Hubert M, Bishop E, Saunders N, Robinot R, Grzelak L, et al. SARS-CoV-2 Alpha, Beta and Delta variants display enhanced Spike-mediated Syncytia Formation. EMBO J. 2021:e108944. Epub 2021/10/04. doi: 10.15252/embj.2021108944. PubMed PMID: 34601723.

45. Schoggins JW. Interferon-stimulated genes: roles in viral pathogenesis. Curr Opin Virol. 2014;6:40–6. Epub 20140405. doi: 10.1016/j.coviro.2014.03.006. PubMed PMID: 24713352; PubMed Central PMCID: PMCPMC4077717.

46. Planas D, Veyer D, Baidaliuk A, Staropoli I, Guivel-Benhassine F, Rajah MM, et al. Reduced sensitivity of SARS-CoV-2 variant Delta to antibody neutralization. Nature. 2021;596(7871):276–80. Epub 2021/07/09. doi: 10.1038/s41586-021-03777-9. PubMed PMID: 34237773.

47. Mahase E. Covid-19: Novavax vaccine efficacy is 86% against UK variant and 60% against South African variant. BMJ. 2021;372:296. Epub 2021/02/03. doi: 10.1136/bmj.n296. PubMed PMID: 33526412.

48. Shen X, Tang H, McDanal C, Wagh K, Fischer W, Theiler J, et al. SARS-CoV-2 variant B.1.1.7 is susceptible to neutralizing antibodies elicited by ancestral spike vaccines. Cell Host Microbe. 2021;29(4):529–39 e3. Epub 2021/03/12. doi: 10.1016/j.chom.2021.03.002. PubMed PMID: 33705729; PubMed Central PMCID: PMCPMC7934674.

49. Kemp SA, Collier DA, Datir RP, Ferreira I, Gayed S, Jahun A, et al. SARS-CoV-2 evolution during treatment of chronic infection. Nature. 2021;592(7853):277–82. Epub 20210205. doi: 10.1038/s41586-021-03291-y. PubMed PMID: 33545711; PubMed Central PMCID: PMCPMC7610568.

50. Corey L, Beyrer C, Cohen MS, Michael NL, Bedford T, Rolland M. SARS-CoV-2 Variants in Patients with Immunosuppression. N Engl J Med. 2021;385(6):562–6. Epub 2021/08/05. doi: 10.1056/NEJMsb2104756. PubMed PMID: 34347959; PubMed Central PMCID: PMCPMC8494465.

51. Lupala CS, Ye Y, Chen H, Su XD, Liu H. Mutations on RBD of SARS-CoV-2 Omicron variant result in stronger binding to human ACE2 receptor. Biochem Biophys Res Commun. 2022;590:34–41. Epub 20211224. doi: 10.1016/j.bbrc.2021.12.079. PubMed PMID: 34968782; PubMed Central PMCID: PMCPMC8702632.

52. Mizuki Yamamoto KT, Youko Hirayama, Jun-ichiro Inoue, Yasushi, Gohda KaJ. SARS-CoV-2 Omicron spike H655Y mutation is responsible for enhancement of the endosomal entry pathway and reduction of cell surface entry pathways. bioRxiv. 2022.

53. Yamamoto M, Gohda J, Kobayashi A, Tomita K, Hirayama Y, Koshikawa N, et al. Metalloproteinase-Dependent and TMPRSS2-Independent Cell Surface Entry Pathway of SARS-CoV-2 Requires the Furin Cleavage Site and the S2 Domain of Spike Protein. mBio. 2022:e0051922. Epub 20220616. doi: 10.1128/mbio.00519-22. PubMed PMID: 35708281.

54. Garcia NK, Lee KK. Dynamic Viral Glycoprotein Machines: Approaches for Probing Transient States That Drive Membrane Fusion. Viruses. 2016;8(1). Epub 20160111. doi: 10.3390/v8010015. PubMed PMID: 26761026; PubMed Central PMCID: PMCPMC4728575.

55. Foster TL, Wilson H, Iyer SS, Coss K, Doores K, Smith S, et al. Resistance of Transmitted Founder HIV-1 to IFITM-Mediated Restriction. Cell Host Microbe. 2016;20(4):429–42. Epub 20160915. doi: 10.1016/j.chom.2016.08.006. PubMed PMID: 27640936; PubMed Central PMCID: PMCPMC5075283.

56. Fenton-May AE, Dibben O, Emmerich T, Ding H, Pfafferott K, Aasa-Chapman MM, et al. Relative resistance of HIV-1 founder viruses to control by interferon-alpha. Retrovirology. 2013;10:146. Epub 20131203. doi: 10.1186/1742-4690-10-146. PubMed PMID: 24299076; PubMed Central PMCID: PMCPMC3907080.

57. Cantuti-Castelvetri L, Ojha R, Pedro LD, Djannatian M, Franz J, Kuivanen S, et al. Neuropilin-1 facilitates SARS-CoV-2 cell entry and infectivity. Science. 2020;370(6518):856–60. Epub 20201020. doi: 10.1126/science.abd2985. PubMed PMID: 33082293; PubMed Central PMCID: PMCPMC7857391.

58. Daly JL, Simonetti B, Klein K, Chen KE, Williamson MK, Anton-Plagaro C, et al. Neuropilin-1 is a host factor for SARS-CoV-2 infection. Science. 2020;370(6518):861–5. Epub 20201020. doi: 10.1126/science.abd3072. PubMed PMID: 33082294.

59. Qing E, Kicmal T, Kumar B, Hawkins GM, Timm E, Perlman S, et al. Dynamics of SARS-CoV-2 Spike Proteins in Cell Entry: Control Elements in the Amino-Terminal Domains. mBio. 2021;12(4):e0159021. Epub 20210803. doi: 10.1128/mBio.01590-21. PubMed PMID: 34340537; PubMed Central PMCID: PMCPMC8406164.

60. Braun E, Hotter D, Koepke L, Zech F, Gross R, Sparrer KMJ, et al. Guanylate-Binding Proteins 2 and 5 Exert Broad Antiviral Activity by Inhibiting Furin-Mediated Processing of Viral Envelope Proteins. Cell Rep. 2019;27(7):2092–104 e10. doi: 10.1016/j.celrep.2019.04.063. PubMed PMID: 31091448.

61. Tartour K, Appourchaux R, Gaillard J, Nguyen XN, Durand S, Turpin J, et al. IFITM proteins are incorporated onto HIV-1 virion particles and negatively imprint their infectivity. Retrovirology. 2014;11:103. Epub 2014/11/26. doi: 10.1186/s12977-014-0103-y. PubMed PMID: 25422070; PubMed Central PMCID: PMCPMC4251951.

62. Brown JC, Goldhill DH, Zhou J, Peacock TP, Frise R, Goonawardane N, et al. Increased transmission of SARS-CoV-2 lineage B.1.1.7 (VOC 2020212/01) is not accounted for by a replicative advantage in primary airway cells or antibody escape. bioRxiv. 2021:2021.02.24.432576. doi: 10.1101/2021.02.24.432576.

63. Thi Nhu Thao T, Labroussaa F, Ebert N, V’Kovski P, Stalder H, Portmann J, et al. Rapid reconstruction of SARS-CoV-2 using a synthetic genomics platform. Nature. 2020;582(7813):561–5. Epub 2020/05/05. doi: 10.1038/s41586-020-2294-9. PubMed PMID: 32365353.

64. Gaida A, Becker MM, Schmid CD, Buhlmann T, Louis EJ, Beck HP. Cloning of the repertoire of individual Plasmodium falciparum var genes using transformation associated recombination (TAR). PLoS One. 2011;6(3):e17782. Epub 2011/03/17. doi: 10.1371/journal.pone.0017782. PubMed PMID: 21408186; PubMed Central PMCID: PMCPMC3049791.

65. Rihn SJ, Merits A, Bakshi S, Turnbull ML, Wickenhagen A, Alexander AJT, et al. A plasmid DNA-launched SARS-CoV-2 reverse genetics system and coronavirus toolkit for COVID-19 research. PLoS Biol. 2021;19(2):e3001091. Epub 2021/02/26. doi: 10.1371/journal.pbio.3001091. PubMed PMID: 33630831; PubMed Central PMCID: PMCPMC7906417.

66. da Silva Filipe A, Shepherd JG, Williams T, Hughes J, Aranday-Cortes E, Asamaphan P, et al. Genomic epidemiology reveals multiple introductions of SARS-CoV-2 from mainland Europe into Scotland. Nat Microbiol. 2021;6(1):112–22. Epub 2020/12/23. doi: 10.1038/s41564-020-00838-z. PubMed PMID: 33349681.

67. Pickering S, Batra R, Merrick B, Snell LB, Nebbia G, Douthwaite S, et al. Comparative performance of SARS-CoV-2 lateral flow antigen tests and association with detection of infectious virus in clinical specimens: a single-centre laboratory evaluation study. Lancet Microbe. 2021;2(9):e461–e71. Epub 2021/07/07. doi: 10.1016/S2666-5247(21)00143-9. PubMed PMID: 34226893; PubMed Central PMCID: PMCPMC8245061.

